# CryoEM Structure of *Drosophila* Flight Muscle Thick Filaments at 7Å Resolution

**DOI:** 10.1101/2020.06.05.136580

**Authors:** Nadia Daneshparvar, Dianne W. Taylor, Thomas S. O’Leary, Hamidreza Rahmani, Fatemeh Abbasi Yeganeh, Michael J. Previs, Kenneth A. Taylor

## Abstract

Striated muscle thick filaments are composed of myosin II and several non-myosin proteins. Myosin II’s long α-helical coiled-coil tail forms the dense protein backbone of filaments while its N-terminal globular head containing the catalytic and actin binding activities extends outward from the backbone. Here we report the structure of thick filaments of the flight muscle of the fruit fly *Drosophila melanogaster* at 7 Å resolution. Its myosin tails are arranged in curved molecular crystalline layers identical to flight muscles of the giant waterbug *Lethocerus indicus.* Four non-myosin densities are observed, three of which correspond to ones found in *Lethocerus;* one new density, possibly stretchin-Mlck, is found on the backbone outer surface. Surprisingly, the myosin heads are disordered rather than ordered along the filament backbone. Our results show striking myosin tail similarity within flight muscle filaments of two insect orders separated by several hundred million years of evolution.

**Significance Statement:** Myosin thick filaments are one of striated muscle’s key structures, but also one of its least understood. A key question is how the myosin a-helical coiled-coil tail is arranged in the backbone. At 7Å resolution, sufficient to resolve individual a-helices, the myosin tail arrangement in thick filaments from the flight muscle of the fruit fly *Drosophila melanogaster* is strikingly similar to the myosin tail arrangement in flight muscles of the giant waterbug *Lethocerus indicus*. Nearly every other thick filament feature is different. *Drosophila* and *Lethocerus* evolved separately >245 million years ago suggesting myosin tail packing into curved molecular crystalline layers forms a highly conserved thick filament building block and different properties are obtained by alterations in non-myosin proteins.

## Introduction

Sarcomeres of striated muscle are composed of four basic components: bipolar, myosin-containing thick filaments; polar, actin-containing thin filaments; a Z-disk which cross-links antiparallel actin filaments into a bipolar structure; and a connecting filament to link the thick filaments to the Z-disk. Of these four elements, the thin filaments are better characterized than the others. Here we are concerned with the myosin-containing thick filaments, the least characterized component structurally.

Molecules of myosin II, the only filament forming myosin (Foth et al., 2006), are heterohexamers consisting of a pair of identical heavy chains of ~2,000 residues and two pairs of light chains (Fig. 1A,B), dubbed essential and regulatory. Myosin’s head comprises the N-terminal ~850 residues plus one of each light chain, the remaining ~1150 residues form a continuous a-helical coiled-coil tail. Myosin heads in thick filaments of relaxed flight muscles of the large waterbug *Lethocerus indicus* extend outwards in intervals of 145 Å (Fig. 1C) giving the appearance of a ring encircling the filament backbone, a structure dubbed a crown.

**Figure 1.**
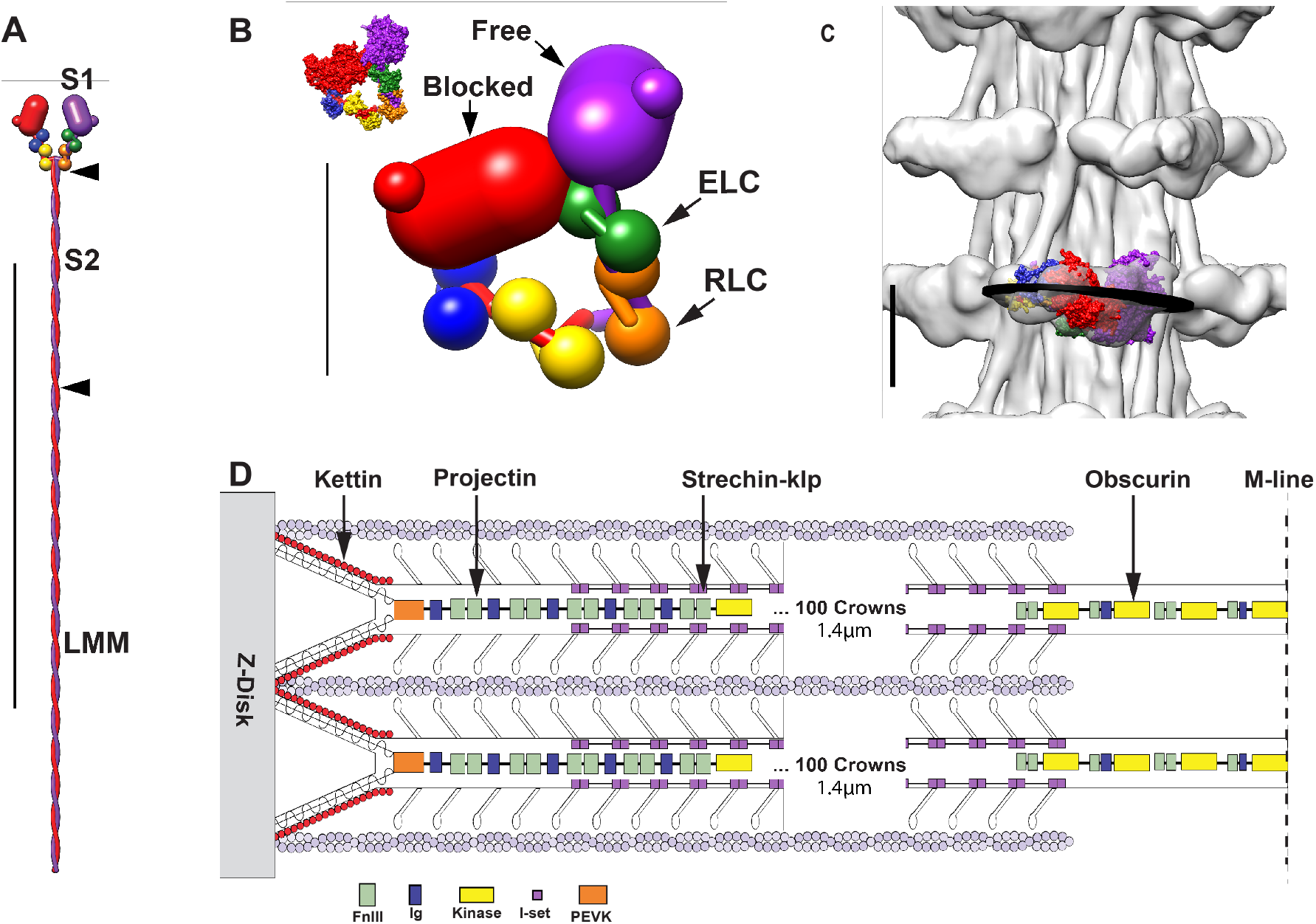
Myosin filament features. (A) Diagram of a myosin molecule with two equivalent heads and an a-helical coiled-coil tail. Proteolysis at two sites (arrowheads) fragments the molecule into two separate heads (S1) and two tail segments [S2 and LMM (light meromyosin)]. Vertical line represents 1000 Å in panel A and 100 Å in B and C. (B) The Interacting Heads Motif (IHM). In the IHM, the two heads are not equivalent. Instead, the actin-binding domain of one head (blocked) contacts the adjacent head (free) whose actin binding domain is not blocked. The inset shows the space-filling structure of PDB 1I84 (Wendt et al., 2001). In filaments, the free head is usually juxtaposed to the thick filament backbone effectively preventing it from binding actin in the relaxed state. (C) The IHM placed within the *Lethocerus* thick filament reconstruction at 20 Å resolution. The black disk approximates the orientation of a best plane drawn through the IHM. This orientation is unique in striated muscle. (D) Schematic diagram showing the relative placement of the giant proteins, kettin, projectin, obscurin and stretchin-klp within a sarcomere. Projectin binds mostly at the filament tip; obscurin to the M-band (bare zone) and kettin to the thin filament and projectin. Stretchin-klp binds along the main shaft of the thick filament but not in the bare zone or at the filament tips. Coloring scheme in A-C – blocked head: heavy chain, red; essential light chain (ELC), blue; regulatory light chain (RLC), yellow. Free head: heavy chain, purple; ELC, green; RLC, orange. Scale bar in A is 1000 Å, in B and C, 100 Å. Coloring scheme for D – Fn3 domains are light green; Ig domains, blue; Ig domains of stretchin-klp, pink; kettin; red and obscurin kinase domains, yellow. D Adapted from (Bullard et al., 2005). A-C from (Hu et al., 2016).

In an active muscle generating tension, individual myosin heads act as independent force generators (Huxley, 1974) and are disordered generally except when attached to actin. In relaxed muscle, in which state the muscle is easily extended because actin-myosin interactions are inhibited, myosin heads become ordered (Huxley and Brown, 1967) though details of their ordered arrangement were obscure for many years. In 2001, the relaxed (inhibited) myosin head conformation of smooth muscle myosin II was visualized (Wendt et al., 2001) and later dubbed an Interacting Heads Motif (IHM; Fig. 1B). All relaxed thick filament structures reported since that resolve individual myosin heads show this same conformation (Craig, 2017). High asymmetry characterizes the head-head interaction of the IHM with the actin binding surface of one head, the blocked head, juxtaposed to the side of the other head, the free head, so named because its actin binding interface is not blocked. In most relaxed thick filaments, the IHM lies roughly tangential to the filament backbone with the free head actin-binding surface facing the backbone, thus preventing actin binding by both heads via different mechanisms. However, in the giant waterbug *Lethocerus* the IHM lies perpendicular to the backbone (Fig. 1C) with the free head bound to the filament backbone by a different mechanism producing an orientation that is so-far unique in striated muscles (Hu et al., 2016). With few exceptions which occur among primitive single cell organisms, ability to form the IHM is nearly ubiquitous for organisms expressing myosin II (Jung et al., 2008, Lee et al., 2018).

While the head is critical for actomyosin motion, myosin’s ~1600 Å long tail is integral to filament formation (Fig. 1A). Using proteolysis in high salt solutions where myosin is soluble, the tail cleaves into two domains, the first ~1/3^rd^ comprising subfragment 2 (S2) and the remaining 2/3^rd^ comprising light meromyosin (Fig. 1A). S2 is soluble at physiological ionic strength, but light meromyosin is not suggesting that light meromyosin contains the thick filament assembly activity with the segment of S2 proximal to the myosin heads providing a tether enabling them to search for and bind actin subunits on the thin filament.

Muscle thick filaments from all species contain additional proteins that modulate their activity or that determine their length. In *Drosophila* flight muscle two of these proteins, projectin and kettin form the connecting filament between thick filaments and the Z-disk (Fig. 1D). Both are largely confined to the filament ends (Ayme-Southgate and Southgate, 2006, Lakey et al., 1990). Obscurin is a bare zone binding protein whose length is about sufficient to reach the first crown of myosin heads (Katzemich et al., 2012). Stretchin-klp is distributed along most of the A-band (Patel and Saide, 2005). Three other proteins (not shown in Figure 1), paramyosin and miniparamyosin (Becker et al., 1992), flightin and myofilin (Qiu et al., 2005) are located in the filament core or among the myosin tails.

Myosin tails provide more than a device for thick filament assembly; they are involved in the activation of myosin heads. Vertebrate skeletal muscle can shorten at low tension even with most myosin heads ordered as in relaxed muscle, but at high loads, the myosin heads become disordered (Linari et al., 2015). The effect, which must be manifest by a change of some kind in the thick filament backbone, has been interpreted as mechano-sensing by the thick filaments (Irving, 2017). Of the ~500 myosin mutations known to cause muscle disease in humans, ~40% are located in the tail domain (Colegrave and Peckham, 2014). Myosin tail mutations may result in incorrect assembly of thick filaments, affect the function of correctly assembled thick filaments, or affect stability resulting in increased turnover. The myosin tail amino acid sequence is highly conserved. The *Drosophila* (fruit fly) and *Lethocerus* (large waterbug) flight muscle myosin tail sequences are 88% identical to each other and when compared to human cardiac β-myosin (MYH7), the sequences are 54% identical, 74% similar.

Asynchronous flight muscle, so named because contractions are out of synchrony with the nervous stimulation, is a comparatively recent adaptation in insects (Pringle, 1981). Four insect orders utilize this type of flight muscle: Diptera (flies, including *Drosophila* sp.), Hemiptera (true bugs, including *Lethocerus* sp.), Hymenoptera (bees and wasps) and Coleoptera (beetles). Asynchronous flight muscles have several characteristics in common (Pringle, 1978) among them are: (1) an indirect arrangement with the muscle attaching to the exoskeleton rather than directly to the wings so that contractions occur at the resonant frequency of the thorax, (2) a high degree of order in the arrangement of thick and thin filaments, (3) relatively sparse sarcoplasmic reticulum, (4) T-tubules aligned with the M-band rather than in the middle of the I-bands or with the Z-disk as occurs in vertebrate striated muscle and in synchronous insect flight muscle (Pringle, 1981), (5) a highly developed stretch activation and shortening deactivation mechanism that enables the muscle to contract rapidly at constant but submaximal calcium concentration, (6) thick and thin filaments nearly completely overlapped at rest length with the result that the amount of shortening is quite small, being only a few percent of the sarcomere length. Stretch activation is most refined in asynchronous flight muscle, but is also observed in vertebrate striated muscle, more so in cardiac than in skeletal muscle (Pringle, 1978).

Recently, a 5.5 Å 3-D image of *Lethocerus* flight muscle thick filaments revealed the myosin tail packing in unprecedented detail along with several unexpected features (Hu et al., 2016). Contrary to the generally accepted model for the myosin tail packing into subfilaments (Wray, 1979), the myosin tails were found organized into curved molecular crystalline layers (Squire, 1973), referred to here as ribbons, due to their rather flat and narrow but elongated morphology. Within ribbons myosin tails from adjacent molecules are offset by 3x 145Å, which favors extensive tail contact within ribbons and somewhat less contact between ribbons (Hu et al., 2016) as well as an enhanced matching of regions of complementary charges (McLachlan and Karn, 1982). Intercalated among the myosin tails were four separate densities with the morphology of extended polypeptide chains. Within the center of the filament was a set of rodlike densities, which are most likely paramyosin.

*Lethocerus* sp. have the distinct size advantage over *Drosophila* for structural studies but cannot be genetically manipulated. *Drosophila* sp. flight muscles can be genetically manipulated often without consequence to their laboratory survival and further breeding. *Drosophila* and *Lethocerus* belong to insect orders that diverged by some accounts ~373 million years ago (Misof et al., 2014). Their flight muscles have very different contraction frequencies and sarcomere lengths. Both have several non-myosin proteins in common, but with some differences in size, sequence and quantity. Here we report two nearly identical 7 Å resolution reconstructed images of relaxed thick filaments from two strains of *Drosophila melanogaster,* a wild type and a regulatory light chain mutant that differ significantly from images of *Lethocerus* in the order of the myosin heads and in the non-myosin proteins. Significantly, the myosin tail arrangements are highly similar. The muscle specific changes appear to be orchestrated around the myosin tail arrangement (ribbons) as a basic structure.

## Results

### 4.1. Thick Filament Appearance in Vitreous Ice

Thick filaments were isolated from flight muscles of two *Drosophila* strains, a wild type (WT), W1118, and a strain designated Dmlc2^[Δ2-46; S66A,S67A]^ (Farman et al., 2009) with regulatory light chain mutations S66A and S67A plus deletion of N-terminal residues 1-46. Regulatory light chain phosphorylation is known to disorder the heads (Levine et al., 1995), but would be impossible in the mutant, which made it a useful test of the possibility that phosphorylation was the agent of myosin head disorder.

Thick filaments of *Drosophila* flight muscle have a length of 3.2 μm (Gasek et al., 2016). Images recorded at a magnification of 18,000x on a DE-64 camera showed almost the entire 2 μm diameter hole in the support film (Fig. 2A,B), facilitating bare-zone and filament tip identification from which filament polarity could be determined *a priori* (Fig. S1A). When the distribution of filament segments included in the reconstruction at the beginning and at the finish is displayed as a histogram, the vast majority come from the region between crowns 0 and 60 (Fig. S1B). The large number of segments including crown 0 is indicative of the frequency with which bare zones were visible.

**Figure 2.**
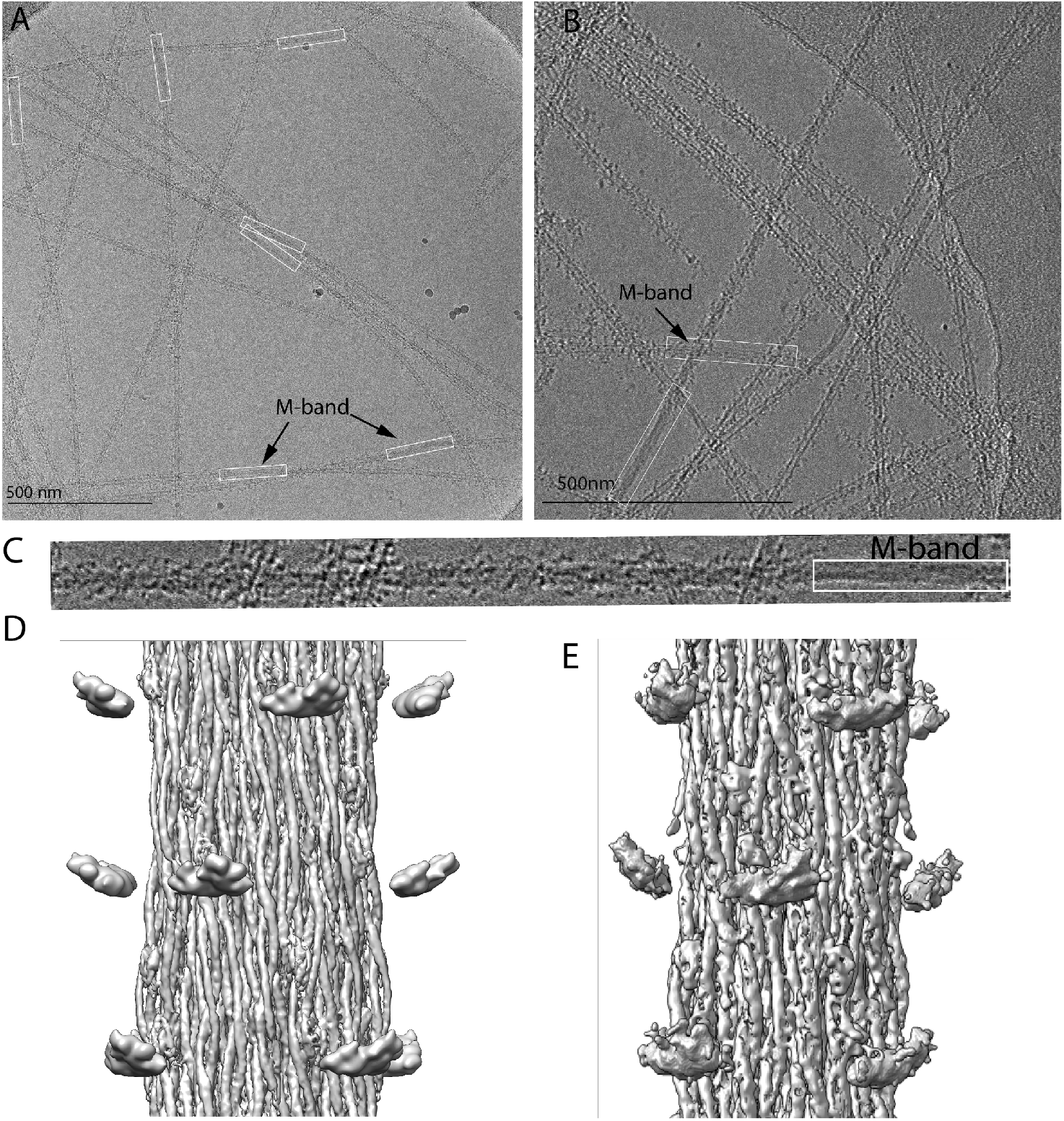
Typical images showing the bare zone (white boxes) and adjacent A-bands of WT and mutant flies. (A) WT isolated thick filaments. In the adjacent A-bands numerous densities (myosin heads) project from the surface. (B) Thick filament from RLC mutant flies. Images for this set recorded using a Volta phase plate. Note that the density across the diameter of the bare zone is uniform but across the A-band is lighter in the middle suggesting the filaments are hollow. (C) Image of the WT thick filament showing the disorder of the myosin heads and the smoothness of the bare zone, outlined in the white box. (D) WT reconstruction after imposing helical symmetry and application of local deblur. (E) Mutant reconstruction after similar treatment. Images in panels D and E low pass filtered to 7 Å resolution for the backbone and 40 Å resolution to display the floating head densities.

Micrographs of frozen hydrated filaments isolated from both strains, generally appeared straight, but bent or broke usually at the bare zone in the filament center (Fig. 2A,B; see also Fig. S1A). Filament density was uniform across the bare zone, but on either end appeared hollow all the way to the filament tip. Images from both sets of thick filaments showed disorder in the myosin heads (Figs. 2A,B) in contrast to the distinct crown structure seen with *Lethocerus* thick filaments (Hu et al., 2016). The head disorder is also apparent at higher magnification (Fig. 2C). After computing the wild type reconstruction we obtained the Dmlc2^[Δ2-46; S66A,S67A]^ flies at which point we also had access to and used a Volta Phase Plate for recording its image data.

The wild type reconstruction showed little detail in the myosin heads, which appeared as floating densities with no visible S2 connection to the backbone (Fig. 2D). Despite enhancement of the filament contrast by the phase plate (Fig. 2B), no significant improvement in myosin head detail was obtained (Fig. 2E).

### 4.2. Reconstruction

Very little density recognizable as myosin heads was visible in both the wild type and Dmlc2^[Δ2-46; S66A,S67A]^ reconstructions. However, resolution of the filament backbone was ~7 Å (Figs. S2,S3) revealing the myosin tail a-helices and non-myosin proteins (Fig. 2D,E). Both reconstructions are nearly indistinguishable and can be described adequately using just the wild type result. Both wild type and mutant had very similar helical angles of 33.86° and 33.92° respectively. Slightly more subfragment-2 is visible in the Dmlc2^[Δ2-46; S66A,S67A]^ reconstruction.

Our *Drosophila* thick filaments have an outer diameter of ~180 Å, slightly more than the ~170 Å predicted for a four stranded filament (Squire, 1973), but slightly less than the 190 Å observed for *Lethocerus.* A 3-stranded thick filament has a predicted backbone diameter of ~155 Å (Squire, 1973). We computed reconstructions from both the wild type and mutant using threerotational symmetries: 2-fold (C2), 3-fold (C3) and 4-fold (C4). The C2 reconstruction was very similar to the C4 but of lower resolution due to less averaging with just C2 imposed; with C3 symmetry imposed, the reconstruction was distinctly different and of lower quality (Fig. S4). Imposing C3 symmetry produced a reconstruction with three robust densities running along the thick filament axis with comparatively flimsy connections very different from the generally uniform cross section density found in other 3-fold symmetric thick filaments at comparable resolution. cisTEM (Grant et al., 2018) was used to compute the final C4 reconstruction. Helical parameters were then determined and imposed using Relion (Scheres, 2012), which produced a smoother map. Lastly the map was sharpened using local DeBlur (Ramirez-Aportela et al., 2020). All these maps are the same in terms of resolution (Figs. S2,S3), but the sharpened map better separates the coiled-coil a-helices and non-myosin proteins.

Extensive averaging of multiple filaments and multiple repeating segments of each filament occurs in the reconstruction process. Consequently, structural elements that do not follow the myosin symmetry, that follow it but are present in less than equimolar amounts, or that are conformationally heterogeneous will not show up in the reconstruction or will only be visualized at a lower contour threshold. Those elements that appear in the reconstruction at the same contour threshold as the myosin tails are present at close to equimolar amounts with respect to myosin and are comparatively conformationally homogeneous.

### 4.3. Myosin Heads are Disordered

Well resolved myosin heads in a modified position of the interacting heads motif characterized the relaxed *Lethocerus* thick filament (Fig. 1C). Surprisingly, no density resembling the interacting heads motif was visible in either *Drosophila* reconstruction even when the map is low pass filtered to 40Å (Fig. 2D,E). Instead four densities occur in the expected axial position of myosin heads but unconnected to the filament backbone and comparatively small and shapeless (Fig. 2D,E), which is characteristic of a disordered structure. Isolation procedures used for *Drosophila* and *Lethocerus* thick filaments were virtually identical (Hu et al., 2016) except that the *Drosophila* thick filaments were made from either fresh tissue, or following brief storage at – 80°C in glycerol buffer. Light chain phosphorylation, which typically disorders the myosin heads (Levine et al., 1996), cannot be the cause because the mutant, which cannot be phosphorylated, also showed disordered heads. The putative myosin head density is axially aligned with the N-terminus of the visible, ordered part of the myosin tail, the proximal S2. In *Lethocerus* thick filaments, the S2 extends ~110 Å from this point to connect the head-tail junction at the next Z-ward crown (Hu et al., 2016); it is this connection that is not visible in *Drosophila.*

### 4.4. Myosin Tails are Arranged in Ribbons

After extending the reconstruction to a length of 12 crowns, a single myosin tail could be segmented from the map (Fig. 3A). Continuous density corresponding to the individual myosin tail a-helices is visible for a distance of 10 crowns; the 11^th^ crown of myosin tail would contain the disordered proximal S2. Myosin tails are arranged within an annulus of outer diameter 180 Å surrounding a hollow core of diameter 70 Å, which is distinct from *Lethocerus* thick filaments where the central core had eight densities believed to be paramyosin. Like the *Lethocerus* thick filament (Hu et al., 2016), the *Drosophila* myosin tail extends from its N-terminus on the outside of the tail annulus through to its C-terminus on the inside giving an outward tilt of 1.8° relative to the filament axis.

**Figure 3.**
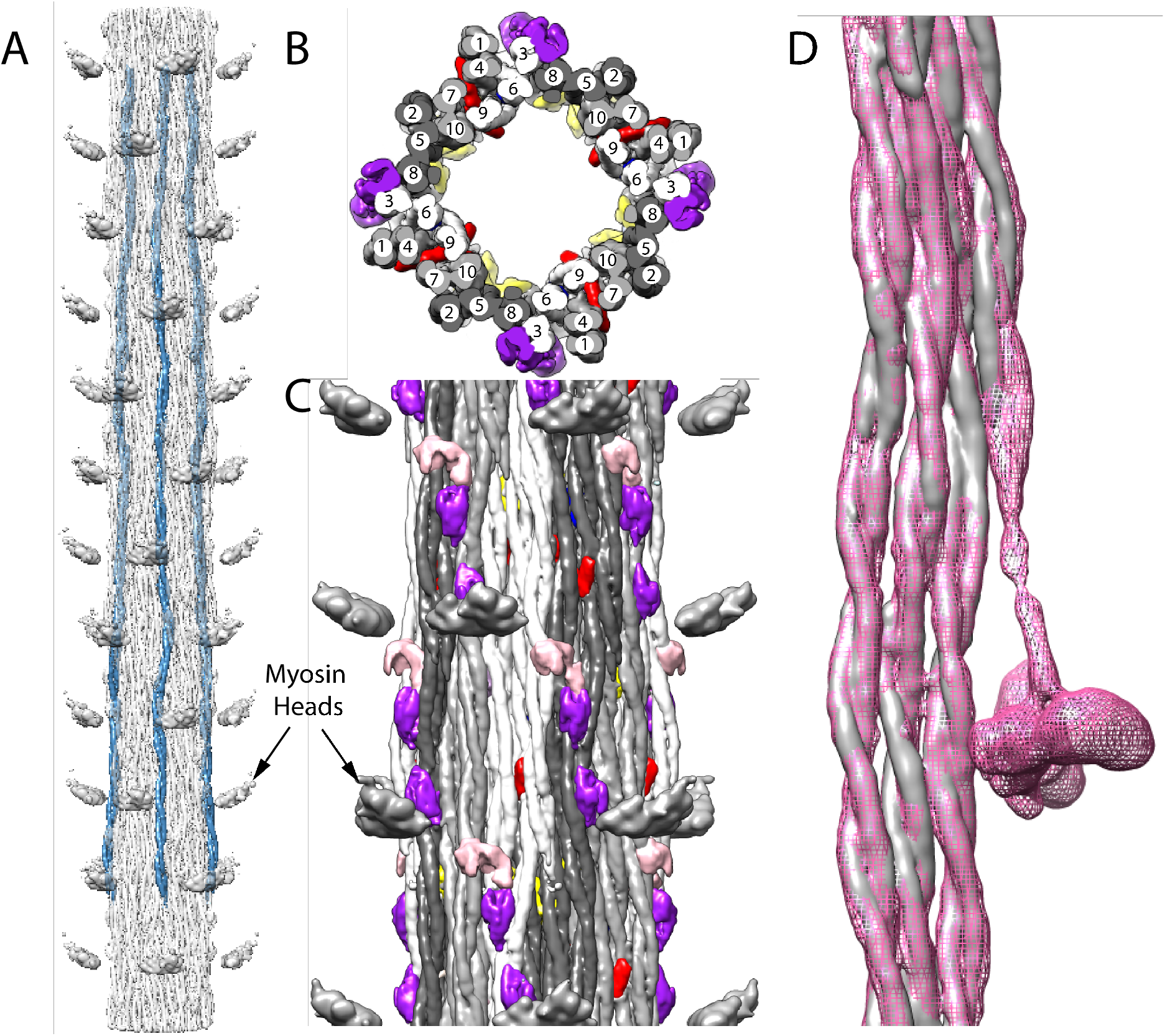
Arrangement of myosin and non-myosin proteins. (A) Four symmetrically placed myosin tails (blue) segmented from a reconstruction extended 12 crowns. As in *Lethocerus* thick filaments, myosin tails run mostly parallel to the filament axis with a slight tilt inwards towards the C-terminus. (B) View looking down the filament axis showing the “curved molecular crystalline layers” (ribbons) (Squire, 1973). Ten myosin tails in each of the four asymmetric units, arising from the 4-fold symmetry, are numbered sequentially according to their 145Å axial offsets. Each ribbon consists of myosin tails offset axially by 3 crowns, i.e. 1, 4, 7, 10; 2, 5, 8; 3, 6, 9. Ribbons are colored white, light and dark gray. (C) Longitudinal view from the outside. Non-myosin proteins flightin (red) and myofilin (yellow) are embedded among the myosin tails. A third, possibly stretchin-klp (purple and pink) is on the outer surface. (D) Myosin tails of *Drosophila* (gray) and *Lethocerus* (pink mesh) superimposed as a ribbon. With the exception of the proximal S2 of *Lethocerus,* which bends to the left, the same feature in *Drosophila* is mostly disordered but what is visible appears to follow a straight trajectory. Note that the density threshold for the *Lethocerus* reconstruction was chosen to be the minimum that would show the position of the free head without blurring the proximal S2.

Evidence for the validity of the ribbon arrangement of myosin tails is provided by the myosin tail packing. A transverse slab through the reconstruction always shows 40 myosin tails due to the 4-fold symmetry of the filament and the 10 crown length of the myosin tail. The proximal S2 extends outside of the myosin tail annulus and is not involved in myosin tail packing within the backbone. These 40 tails are arranged in 12 ribbons. By symmetry, a 3-crown length of a ribbon contains a complete myosin molecule, cut into three 3-crown lengths and one extra crown length. Thus, a transverse slab through a ribbon shows a width of three myosin tails 2/3^rds^ of the time, and 1/3^rd^ of the time a fourth myosin tail. Segmentation of all the myosin tails revealed a ribbon packing very similar to that found for *Lethocerus* (Fig. 3B,C) with myosin tails offset by 3 crowns, the optimal offset predicted by the amino acid sequence (McLachlan and Karn, 1982). When a segmented *Lethocerus* ribbon is superimposed on one from *Drosophila*, there is almost complete superposition (Fig. 3D; Supplemental Movie 5) with the exception of the disordered *Drosophila* proximal S2.

### 4.5. Identification of Non-myosin Proteins

After segmenting the reconstruction, four groups (following the imposed 4-fold symmetry) each with three non-myosin densities were visible among the myosin tails (Figs. 3B,C, 4A-E). On the backbone surface were four additional groups each with three non-myosin densities (Fig. 4F-H). The major reported non-myosin, thick filament associated proteins in *Drosophila* flight muscle are flightin (Vigoreaux et al., 1993), myofilin (Qiu et al., 2005), stretchin (Champagne et al., 2000, Patel and Saide, 2005), paramyosin, miniparamyosin (Maroto et al., 1996), kettin (Lakey et al., 1993), projectin (Hu et al., 1990), and obscurin (Katzemich et al., 2012). Can any of these non-myosin proteins account for these densities?

**Figure 4.**
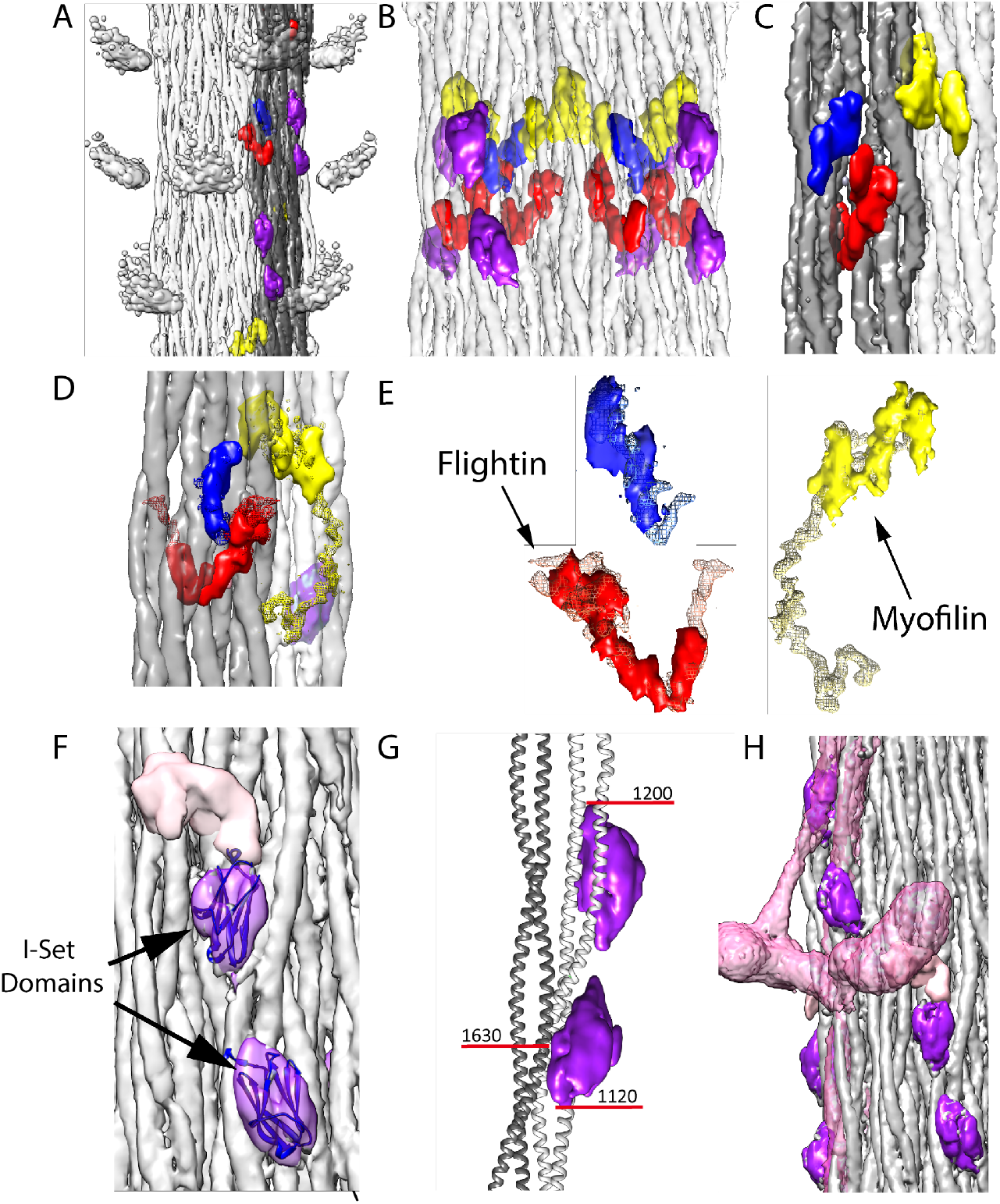
Non-myosin proteins of the *Drosophila* thick filament reconstruction. Coloring scheme same as Figure 3. One ribbon is colored gray. Myosin tails in panels A, B, and D are at 50% transparency. (A) View from the outside showing the myosin head density in relationship to the non-myosin proteins. (B) View from the outside at higher magnification looking through the myosin tails. The putative stretchin-klp densities (purple and pink) of *Drosophila*, which are not seen in *Lethocerus*, are found only on the outside of the thick filament backbone. (C) View from the inside looking out. Flightin (red) extends through the gray ribbon, whereas myofilin (yellow) is at the edge of the ribbon and the blue protein positioned on its surface. (D) The blue density binds the inner surface of the gray ribbon; the yellow (myofilin) density binds between the gray and white ribbon; the red density penetrates the gray ribbon. (E) Red, blue and yellow non-myosin densities of *Drosophila* superimposed on the corresponding feature of *Lethocerus* displayed as mesh. (F) View from the outside showing the putative stretchin-klp density. An atomic model of an Iset domain has been built into two of the densities. The third (pink) is shown without an atomic model. Threshold for I-set/tail is 4.65 and threshold for stretchin linker is 1.00. (G) The paired Ig-like densities shown on a ribbon model of the myosin tail. (H) The putative stretchin-klp densities superimposed on a *Lethocerus* thick filament reconstruction where they appear to pass under the free head and S2. Note that the density threshold for the *Lethocerus* reconstruction was chosen to be the minimum that would show the position of the free head without blurring the proximal S2.

The molar abundances of non-myosin proteins in both myofibrils and isolated thick filaments were quantified by label-free mass spectrometry analyses. Isolation protocols for both myofibrils and thick filaments were nearly identical to those used in the cryoEM studies except for an additional centrifugation step to separate filaments from proteins in the supernatant. For mass spectrometry, samples were digested to peptides with trypsin. The resultant peptides were separated by ultra-high-pressure liquid chromatography and directly analyzed by electrospray ionization mass spectrometry (Methods). The relative abundance of myosin to each non-myosin protein was determined from the liquid chromatography peak areas of the three most abundant peptides from the myosin heavy chain, essential and regulatory light chains, and non-myosin proteins (O’Leary et al., 2019). These relative abundances were divided by 2 (except for paramyosin which is a dimer) to determine the average number of double-headed myosin molecules per non-myosin thick filament associated protein (Table 1). Although myofibrils were present in sufficient quantity for measurement in triplicate, sufficient isolated filaments were available for only a single measurement.

**Table 1.** Summary of Mass Spectrometry Results

| Gene | Common name | Accession No. | Molecular Weight (Da) | Myosin molecules per protein molecule (myofibril) | Myosin molecules per protein molecule (filaments<sup>a</sup>) |
| --- | --- | --- | --- | --- | --- |
| fln | Flightin | P35554 | 20,656 | 1.3 ± 0.4 | 1.5 ± 0.5 |
| Mf | Myofilin | Q9VFC7; C1C553 | 41,667 | 2.0 ± 0.7 | 2.0 ± 0.6 |
| strn-mlck | Stretchin-Mlck | A1ZA73 | 215,065 | 3.4 ± 1.1 | 7.9 ± 2.5 |
| prm | Paramyosin | P35415 | 102,338 | 23.5 ± 7.9 | 30.5 ± 9.6 |
| prm | Mini-paramyosin | P35416; M9NDM6 | 74,277 | 9.7 ± 3.3 | 14.7 ± 4.6 |
| bt | Projectin | L0MN91 | 992,527 | 30 ± 10 | Not detected |
| sls | Titin/Kettin | M9PEA0 | 2,102,814 | 34 ± 12 | Not detected |
| unc-89 | Obscurin | A8DYP0 | 475,000 | 42 ± 14 | 62 ± 20 |

<sup>a</sup>Sufficient specimen for only a single measurement. Uncertainties were determined from variability between the relative ratios generated using peptides from the myosin heavy chain, essential and regulatory light chains.

Flightin, myofilin, stretchin, paramyosin and miniparamyosin were the most abundant non-myosin thick filament associated proteins in the *Drosophila* myofibrils and the most likely to be seen in the reconstruction. Their abundance was consistent with previous biochemical observations (Beinbrech et al., 1985, Ayer and Vigoreaux, 2003, Qiu et al., 2005). Kettin, projectin, and obscurin were present at lower stoichiometries in the myofibril samples. Kettin and projectin were largely absent in the thick filament samples. Kettin in particular is susceptible to calpain cleavage (Lakey et al., 1993) which may be the agent of its loss. Kettin and projectin are mostly at the filament tips where very few segments were selected for the reconstruction (Fig. S1B). They thus make a negligent contribution to the reconstruction. Obscurin is a M-band protein with little extension beyond the first crown of myosin heads. Although present, its contribution is heavily diluted by the overweighting of segments beyond where it could contribute.

Three non-myosin densities (red, yellow and blue in Figs. 3B,C; 4A-E; Supplemental Movies 1-3) are similar to ones observed in *Lethocerus* thick filaments. One density (red) penetrates a ribbon, and corresponds in location and shape to the putative flightin in *Lethocerus* thick filaments (Fig. 5A,B) (Hu et al., 2016). Flightin’s nearly 1:1 stoichiometry with myosin (Table 1) (Ayer and Vigoreaux, 2003, Qiu et al., 2005) is consistent with this assignment. *Drosophila’s* putative flightin density does not extend as far outside the filament backbone as in *Lethocerus* (Fig. 4E), where it made a contact with the proximal S2, which is disordered in *Drosophila,* thus disordering any flightin density associated with it. A second non-myosin density held in common (yellow) was tentatively identified as myofilin in *Lethocerus* (Hu et al., 2016). Myofilin has a slightly lower stoichiometry than flightin with respect to myosin molecules (Table 1) and in *Drosophila* is significantly smaller than in *Lethocerus* (Fig. 4E). The *Lethocerus* putative myofilin density had a domain close enough to paramyosin to suggest an interaction; it is this domain that is visible here even though paramyosin is not. Both putative myofilin and flightin densities in *Drosophila* are too small to contain a full molecule. A third non-myosin density (blue in Figs. 4A-E; Supplemental Movie 3), which may be a portion of either flightin or myofilin, linked by a disordered peptide, remains unidentified in both species. Its size and shape is nearly identical to that of *Lethocerus.*

Calpain clevage can be detected by mass spectroscopy as the absence of peptides following digestion. The N-terminal 64 residues of flightin were not detected in the filament preparations but were detected in the myofibril digests. We detected no calpain cleavage of myofilin in the filament preparations. Note that previous reports of calpain cleavage of flightin and myofilin in *Lethocerus* flight muscle were obtained from Z-disk preparations, not from thick filament preparations (Bullard et al., 1990). Calpain cleavage of stretchin-klp was also detected (see below).

Three additional densities not observed in *Lethocerus* (purple and pink in Figs. 3B,C, 4F-H; Supplemental Movies 1-4) were observed on the outside surface of the *Drosophila* thick filament ‘ backbone. The two purple densities were similar but not identical. At a somewhat lower contour threshold the pink density appears, suggesting that another feature is present closely linked to the purple densities but with lower occupancy or greater heterogeneity. Based on the proteomic data, these three additional densities appear to be from stretchin-mlck’s Ig-like domains and linkers.

In both the filaments and the myofibrils, 15 peptide fragments from the trypsin digests are found covering residues 128-1843 of A1ZA73. A shorter transcript of stretchin-mlck, pC1, has been reported (Patel and Saide, 2005) consisting of 711 residues, contained entirely within A1ZA73, residues 232-940 with the exception two serines added at positions 1 and 2. The first A1ZA73 fragment overlapping pC1 occurs at residues 458-465 so no observed peptide contained the N-terminal serines of pC1. The full length A1ZA73 appears to be expressed in both our myofibril and filament preparations.

Stretchin-mlck is a large gene from which seven potential transcripts have been predicted (Champagne et al., 2000). One of these is a kettin-like protein, referred to as stretchin-klp (Patel and Saide, 2005), which is the form detected in our proteomics. Stretchin-klp (Uniprot, isoform R, A1ZA73) consists of an unstructured N-terminal 455 residues followed by five repeats of Ig-like – short linker – Ig-like – long linker. Short linkers are 11-27 residues long and long linkers are 61-174 residues long. Full length stretchin-klp is detected in myofibrils, but in our filament preparations, the first 455 residues are not detected, presumably clipped by calpain. Assuming that the variable long linkers form different folded structures, a sequence of five 3-domain repeats can be formed. Some servers, e.g. PubMed, predict six three domain repeats. Regardless of five or six repeats, the predicted pattern is a pair of Ig-like repeats separated by a short linker and linked to successive pairs by a long linker. Because each repeat corresponds to one myosin dimer, the expected ratio of myosin:stretchin-klp is 5 (or 6), close to the ratio found in the myofibril preparation, 3.4 ± 1.1 and in the filaments 7.9 ± 2.5.

Each purple density is fit well by the atomic structure of an I-set domain of myosin binding protein C (PDB 2YXM; Fig. 4D), which is a type of Ig domain found in many striated muscle proteins and predicted for stretchin-klp (Supplemental Movie 3). Both stretchin-klp Ig domains are positioned on or near the Skip 1 region of one myosin tail on the backbone surface (Fig. 4E; Supplemental Movie 6), identifiable by the parallel (uncoiled) a-helices characteristic of the Skip 1 region (Taylor et al., 2015). Using the distance along the myosin tail density as a measure of residue number, the contact site on one myosin tail for both Ig-like domains falls between residues 1120-1200 (Fig. 4E). One of the Ig-like domains appears to contact a myosin tail from an adjoining ribbon centered approximately on residue 1630. Chains of stretchin-klp molecules could follow either left-handed or right-handed helical tracks with the shorter distance between three-domain repeats being along left-handed tracks. (Fig. 4F, Fig. S5A).

Was paramyosin present in the thick filament reconstructions? In *Drosophila* myofibrils, the ratio of myosin to paramyosin, 23.5 ± 7.9:1 (Table 1), was lower than the ratio for isolated filaments, 30.5 ±9.6. Both ratios exceed the 15.4:1 ratio determined from *Drosophila* flight muscle (Beinbrech et al., 1985), which is twice the ratio of myosin heavy chain to paramyosin (8.2:1, 7.7:1) for *Lethocerus* thick filaments (Bullard et al., 1973, Levine et al., 1976). Although a putative paramyosin was visible in *Lethocerus* thick filaments, its visualization required that the contour threshold be lowered since it did not appear to follow the myosin helical symmetry. Thus, the lower paramyosin content in *Drosophila* thick filaments could account for its invisibility when the map is contoured at thresholds suitable for visualizing the myosin tails.

### 4.6. Visibility of Myosin Heads

Our *Drosophila* structure shows no well-resolved heads. To find an explanation, we aligned a ribbon segmented from the *Lethocerus* reconstruction (EMD-3301) with our *Drosophila* reconstruction with the result that one of the *Drosophila* putative stretchin-klp domains was positioned beneath the presumptive location of the proximal S2 region and the other one positioned close enough to the free head that it could sterically prevent binding to the filament backbone (Fig. 4H, Fig. S5B,C). Thus, it would seem that the purple and pink densities, which mass spectroscopy indicates are stretchin-klp, are preventing formation of an ordered interacting heads motif similar to that of *Lethocerus*.

## 5. Discussion

Thick filaments from invertebrate striated muscles are ideal specimens for probing the arrangement of myosin molecules as well as accessory proteins, because the filaments are helical in structure over extended lengths. It is no surprise that so far the highest resolution thick filament structures have come from invertebrate thick filaments such as the large water bug *Lethocerus indicus* (Hu et al., 2016) and tarantula (Yang et al., 2016). Invertebrate thick filaments are highly variable between species, offering possibilities for comparative studies.

The primary disadvantage of water bugs and tarantula is that neither comes from a genetic model organism, which inhibits production of new modified strains for hypothesis testing. Many mutations of *Drosophila* flight muscle thick filaments exist. Flight muscle thick filaments from both *Drosophila* and *Lethocerus* possesses several similarities but also had some surprising differences.

### 5.1 Why are Myosin Heads Disordered

Even partially ordered myosin heads could not be obtained using procedures that worked well for *Lethocerus* thick filaments. Within regions from which myosin heads project, the filaments appear well preserved and generally straight. Isolation procedures were the same for both species but were performed on much fresher *Drosophila* muscle. Conceivably prolonged glycerination or calpain treatment resulted in loss of components from *Lethocerus* filaments but it is equally likely that *Lethocerus* does not express a stretchin-klp ortholog.

Early efforts by others to preserve thick filaments from *Lethocerus* in vitreous ice using completely different specimen preparation methods showed well-ordered crowns of myosin heads, but disordered heads with *Drosophila* thick filaments (Menetret et al., 1990). Our methods handle the isolated filaments as little as possible to preserve ordering of the myosin heads. Thus, it seems unlikely that the specimen preparation protocols used here are responsible for the differences.

Regulatory light chain (RLC) phosphorylation is known to disorder myosin heads in vertebrate thick filaments (Levine et al., 1995). However, the same isolation procedures performed on the mutant fly strain Dmlc2^[Δ2-46; S66A,S67A]^, whose RLC phosphorylation sites and N-terminal 46 residues had been removed (Farman et al., 2009) produced no better head ordering than wild type. Thus, RLC phosphorylation was not the cause of the head disorder. Within intact muscle fibers, the Dmlc2^[Δ2-46; S66A,S67A]^ mutant shows less order than wild type, possibly caused by the inability of the myosin heads to bind thin filament via the RLC N-terminal extension (Farman et al., 2009). Absent thin filaments, wild type and mutant isolated thick filaments were indistinguishable.

*Drosophila* filaments have much less paramyosin in their core than *Lethocerus* filaments which might make them more fragile. However, broken filaments are relatively infrequent in both preparations and are usually avoided. Filaments bent at the bare zone are included in the data analysis; those bent in the A-band are not. Although this possibility cannot be excluded at this time, we think low paramyosin content is an unlikely cause.

Do myosin heads in relaxed *Drosophila* thick filaments form an interacting heads motif? If not, they would be the first striated muscle system that does not, as all other relaxed thick filament structures have shown this inactive myosin motif (Craig, 2017, Hu et al., 2016). Relaxed *Lethocerus* thick filaments also have the interacting heads motif, but in an altered orientation that does not involve subfragment 2. Instead of the blocked head binding subfragment 2 and holding the free head actin-binding interface against the filament backbone, free head binding to the filament backbone in *Lethocerus* is independent of the blocked head, which binds only the free head (Hu et al., 2016). *Drosophila* flight muscle myosin in solution forms the interacting heads motif (Lee et al., 2018), but if it forms in filaments, it may be disordered due to amino acid substitutions that eliminate key interactions between myosin heads and myosin tails such as occur with the *Lethocerus* free head. The flight muscle myosin tail sequences of *Lethocerus* and *Drosophila* myosin are 88% identical and 98% similar so this would seem unlikely.

*Drosophila* thick filaments have protein densities on their surface that are absent in *Lethocerus* and which could provide a steric block to free head binding. Stretchin-klp, which consists of a string of 10 (or 12) Ig domains appears to be the most likely candidate. Stretchin-klp has been studied in *Drosophila* flight muscle using antibodies, which showed labeling from a location 0.1 μm from the M-line for a distance of ~1.2 μm (Patel and Saide, 2005), a region that closely approximates the part included in our reconstruction (Fig. S1B). Proteomic analysis indicates that the full-length form is present in the myofibrils, but in the isolated thick filaments, the first 455 residues were missing leaving only the Ig domains and their linkers. Putative stretchin-klp densities follow a left-handed helical path that passes under the location where the free head of the *Lethocerus* interacting heads motif would be positioned in relaxed thick filaments (Figs. 4F,S5; Supplemental Movie 6). Thus, a steric block to free head binding on the thick filament surface as occurs in *Lethocerus* seems the most likely explanation for the lack of myosin head order in *Drosophila*. The blocked head and free head could still interact, but lacking a binding site on the thick filament backbone or S2, the interacting heads motif, if present, would be disordered.

### 5.2. Comparison with *Lethocerus* Thick Filaments

Thick filament backbones of *Lethocerus* and *Drosophila* show both similarities and differences. No density assignable to paramyosin is observed. Density putatively assigned to paramyosin in *Lethocerus* was low compared to myosin and probably does not represent accurately the arrangement of paramyosin molecules (Hu et al., 2016). *Drosophila* has about half the paramyosin per myosin molecule (1:15) as *Lethocerus* (1:7) which would explain its invisibility in the reconstruction (Table 2). *Lethocerus* thick filaments had an additional non-myosin density (colored green in (Hu et al., 2016)) that now appears to be the non-helical myosin C-terminus of the blocked head heavy chain (Rahmani, in preparation) but is possibly disordered in *Drosophila*.

**Table 2.** Comparison of *Lethocerus* and *Drosophila* thick filament proteins

|  |  | *Lethocerus indicus* |  | *Drosophila melanogaster* |  | Reference |
| --- | --- | --- | --- | --- | --- | --- |
| Filament length | | 2.3 $\mu\text{m}$ | | 3.2 $\mu\text{m}$ | | (Reedy, 1967, Gasek et al., 2016) |
| Rotational symmetry |  | C4 |  | C4 |  | (Reedy et al., 1981, Morris et al., 1991); this work) |
| Helical Parameters |  | 145Å, 33.98° |  | 145Å, 33.86° |  | (Hu et al., 2016); this work) |
| Non-Myosin Proteins | Protein | Molecular weight | Ratio to Mhc | Molecular weight | Ratio to Mhc |  |
|  | Flightin | 19 kDa | 1:2 | 20 kDa | 1:2 | (Qiu et al., 2005) |
|  | Myofilin | 30 kDa | 1:2 | 20 kDa | 1:2 | (Qiu et al., 2005) |
|  | Paramyosin | 107 kDa | 1:7<sup>a</sup> | 102 kDa | 1:15 | (Becker et al., 1992, Vinos et al., 1991, Bullard et al., 1973) |
|  | Mini-paramyosin | 62 kDa |  | 55 kDa |  | (Maroto et al., 1996, Becker et al., 1992) |
|  | Projectin | 800 kDa |  | 800-1000 kDa |  | (Ayme-Southgate and Southgate, 2006, Lakey et al., 1990) |
|  | Kettin | 700 kDa |  | 540 kDa |  | (Bullard et al., 2006, Lakey et al., 1993) |
|  | Stretchin-klp |  |  | 225, 231 kDa, |  | (Patel and Saide, 2005) |

<sup>a</sup> *Lethocerus cordofanus* and *Lethocerus maximus*

Three non-myosin densities in *Drosophila* are similar to ones previously found in *Lethocerus* (Hu et al., 2016)*. Drosophila* thick filaments have a V-shaped feature, colored red here and tentatively identified as flightin, that corresponds closely in shape to the similar density seen in *Lethocerus*, including its penetration through a ribbon and a folded globular domain of similar shape and position on the inner surface. Only the part contacting the proximal S2 outside the backbone is missing here either because the proximal S2 itself is disordered or it was clipped by clapain. *Drosophila* and *Lethocerus* flightin are similar in size (Table 2) and have regions of sequence in common, although the entire sequences are only 35% identical (Qiu et al., 2005) compared to 88% identical in the myosin tail. Among the differences are their hydrophobicity, number of isoforms and phosphovariants (11 isoforms in *Drosophila*, 3 in *Lethocerus*; 9 phosphovariants in *Drosophila*, 0 in *Lethocerus*). Hydrophobicity in *Drosophila* flightin is low in contrast to high hydrophobicity in *Lethocerus* (Barton and Vigoreaux, 2006). Flightin is sensitive to calpain cleavage (Bullard et al., 1990) which was also observed in our preparations.

*Lethocerus* thick filaments had a second density, also present in *Drosophila*, tentatively identified as myofilin (yellow in Figs. 3, 4). *Drosophila* myofilin is ~10 kDa smaller than its *Lethocerus* ortholog and also has a smaller density visible in the reconstruction. In *Lethocerus*, the myofilin density has a globular core that might contact paramyosin and an extended domain juxtaposed to the myosin tail annulus. Only the globular core is seen in *Drosophila*; the extended domain is missing (Fig. 4E). The myofilin volume resolved in both *Drosophila* and *Lethocerus* is less than the molecular weight would suggest; other parts remain to be resolved or are disordered.

The blue density here is nearly identical in shape to the similar density in *Lethocerus* (Fig. 5A,B). It is the only non-myosin density that appears nearly identical in both reconstructions, but which has yet to be even tentatively identified, thoug it could be part of either flightin or myofilin.

Similarity would be expected given the high sequence homology in the coiled-coil tail, but seems surprising given the differences observed in other locations of the reconstruction. High sequence homology characterizes the myosin coiled-coil tail domains: >88% identity among the four insect orders having asynchronous, indirect flight muscles, 60-75% identity among tarantula, horseshoe crab and scallop, three species for which thick filament reconstructions have been published, and 50-54% identity between *Drosophila* flight muscle and human striated muscle (Hu et al., 2016). Differences between *Drosophila* and *Lethocerus* flight muscle thick filaments appear in the myosin associated proteins and the myosin head order, not in the myosin tail arrangement.

*Drosophila* and *Lethocerus* flight muscles differ in structure and performance including: (1) *Drosophila* wing beat frequency at 200 Hz is ~5x faster than *Lethocerus* (Dickinson et al., 2005), (2) *Drosophila* flight muscle myosin actin activated ATPase is correspondingly faster than *Lethocerus’* (Swank et al., 2006), (3) *Drosophila* thick filaments are 40% longer than *Lethocerus’* and have half as much paramyosin, (4) adjacent thick filaments in *Drosophila* flight muscle are axially staggered by 145Å/3 to produce a superlattice (Squire et al., 2006) whereas *Lethocerus* flight muscle has no such superlattice (Schmitz et al., 1994), (5) if *Drosophila* forms an interacting heads motif in relaxed filaments, an unanswered question, it is disordered.

### 5.3. Conclusion

A remarkable parallel appears between the consistency of the ribbon as a structural building block of thick filaments and the consistency of the actin subunit as the structural building block of thin filament. Thin filaments across most if not all striated muscles have in common very similar F-actin structures (Galkin et al., 2002), highly conserved due to actin’s high sequence conservation, differing basically by small, though not insignificant, changes in helical angle. They differ mainly in the F-actin binding proteins in particular troponin and tropomyosin, changes in which produce the different functional properties (Bullard and Pastore, 2019). Hemiptera, of which *Lethocerus* is a member, are thought to have appeared in the Devonian period ~373 million years ago (Misof et al., 2014). Diptera of which *Drosophila* is a member appeared later, in the early Triassic, about 245 million years ago. Asynchronous flight muscle is thought to have evolved independently a number of times (Pringle, 1978, Pringle, 1981) so it is not certain that Diptera and Hemiptera had a common ancestor with asynchronous flight muscle. Possibly more remarkable would be the independent evolution to such a similar ribbon arrangement of myosin tails and similar helical angle. Since the Hemiptera and Diptera appeared so long ago, we expect and observe distinct differences in their flight muscle thick filaments. However, the myosin tail structure and arrangement in ribbons are nearly identical implying a strong evolutionary pressure towards maintenance. Other features, such as the associated proteins are very different suggesting adaptation to physiological requirements are affected by the non-myosin proteins. In this context, the myosin head, which can evolve separately from the tail, is a myosin tail associated protein.

Invertebrates appeared long before vertebrates yet even among highly different species as *Placopecten magellanicus* (scallop) and *Drosophila* the sequence identity within the myosin rod is as high as 75%. Vertebrates appeared some 480 – 360 million years ago by some estimates (Sallan et al., 2018) yet vertebrate thick filaments from different species are highly similar in structure at least to the extent that comparative studies have been done. For example, they have similar length, same rotational symmetry, have titin to determine their length, and an axial repeat of 429 Å (3 x 143 Å). It is perhaps worth speculating that the curved molecular crystalline layers (ribbons) first proposed in 1973 (Squire, 1973) may be far more similar to each other across species and muscle types than suggested by the myosin tail sequence identity. X-ray fiber diffraction studies of vertebrate striated muscle showed a clear preference for the ribbon structure over other models based on subfilaments (Chew and Squire, 1995). High amino acid sequence conservation of the myosin tail from many striated muscles, particularly for the charged residues important for filament formation (McLachlan and Karn, 1982) suggest that among the many different kinds of striated muscle myosin filaments, that the ribbons may be nearly identical. Differences among the filaments would be accounted for by small changes in the arrangement of ribbons to produce different rotational and helical symmetries accompanied by significant changes in the non-myosin proteins as adaptations to differing physiological requirements.

## 6. Materials and Methods

Calcium insensitive human plasma gelsolin (residues 25-406) was cloned and overexpressed in *Escherichia coli* BL21 (DE3) strain in Lysogeny Broth (LB) medium. The expression vector was obtained from Dr. Margaret Briggs (Duke University Medical Center). The detailed protocol is included in the Supplemental Information.

### 6.1 Thick Filament Preparation

For a typical preparation, indirect flight muscle is dissected from the thorax of 10 *Drosophia* flies. Typically, flies are immobilized with CO2 and then dissected by removing the heads, abdomen, wings, and legs leaving only the thorax. The vast majority of protein in the thorax is indirect flight muscle. We made no attempt to separate the dorsal longitudinal from the dorsal ventral muscle or any of the nonfibrillar muscles, which are much smaller and present in much lower quantity. Further details are in the Supplemental Methods

### 6.2 Electron Microscopy: grid preparation

Thick filament preparations varied in concentration. Rather than centrifuge them to increase their concentration, we varied the number of drops of suspension applied to the grids, blotting away the excess from the side opposite where the drop was applied. Before proceeding to freezing, the concentration was evaluated by preparing a negatively stained specimen exactly as for freezing except that a final wash in 1% uranyl acetate negative stain was performed, the grids air dried, and examined in a CM120 electron microscope rather than being plunge frozen.

Frozen specimens were prepared on plasma-cleaned 200 mesh Cu Quantifoil R2/1 reticulated carbon grids using the back-blotting technique (Toyoshima, 1989). The concentration of filaments was relatively low in these preparations. By using the back-blotting technique with reticulated carbon grids, the carbon film acts as a sieve to trap several filaments over the 2 μm diameter holes often unobstructed by crossing filaments (Figure 2A,B). The grids were then plunge-frozen in liquid ethane, cooled by liquid nitrogen, using a home-built plunge-freeze device mounted in a 4° cold room.

The grid prepared in this manner have relatively uniform ice thickness with some areas of thicker ice towards the center of the grid square with thinner areas around the perimeter. We picked holes for data collection with medium levels of ice thickness for imaging to avoid as much as possible the possibility that contact with the air-water interface could be the agent of head disorder.

### 6.3 Electron Microscopy: Data Collection

The wild type sample grids were imaged on a FEI Titan Krios transmission electron microscope operated at 300kV. Grids were maintained at liquid nitrogen temperatures at all times after preparation. Images were recorded on a Direct Electron Ltd DE-64 camera operated in integrating mode with 44 frames collected with a total electron dose of 26 e^-^/Å^2^.

Image data were recorded a DE-64 direct electron detector, which has a frame size of 8192 x 8192 pixels. A total of 1510 mages were collected at 18,000x magnification (2.07 Å/pix) with a total exposure of 26 e^-^/Å^2^ and a defocus range of −4 to −6 μm.

The second set of data was collected for the mutant Dmlc2^[Δ2-46; S66A,S67^A] using the Volta phase plate and DE-64 in integration mode, which had recently become available to us. A total of 2002 images were collected at 29000x magnification (1.29 Å/pix) with total dose of 26 e^-^/Å^2^,21 frames, and defocus range of −0.2 to −1.0μm (Fig. 2B).

### 6.4 Data Analysis

After frame alignment and CTF correction, we manually picked filaments using Appion manual picker and used RELION (He and Scheres, 2017) helical extraction to obtain the particle stack. We started with 60,000 segments with a box size of 324 x 324 pixels, which contains approximately four axial repeats of length 145 Å. The 145 Å repeat is generally referred to as a “crown” (Taylor et al., 1984). In addition to a generally low signal to noise ratio, the myosin heads in our thick filaments are disordered, which could throw off the alignment significantly, unless other features within the backbone can define the axial repeat period. Initial 2D classification into 50 classes by Relion was highly inconclusive. The averages were featureless which showed the programs inability to align particles and classify them effectively (Suppl. Fig. S2A). To obtain better definition among the classes, we tested ROME 1.1 (Wu et al., 2017) for 2D classification. ROME 1.1, which uses statistical manifold learning (SML), could not handle a large unbinned data set. Therefore, we binned the stack by 3 (box size of 128). We first ran the utility *rome_map* with 50 classes and started to see features in the backbone even though density corresponding to myosin heads remained unclear. After selecting the classes showing the clearest backbone features, we ran the utility *rome_sml* requesting 300 classes (Fig. S2B). SML turned out to be a powerful classification tool for this data. After removing non-sensical classes ~149,000 segments remained.

Unfortunately, multiple efforts at 3D classification in both RELION and ROME failed. We then imported the best classes obtained from *rome_sml* and reprocessed them using cisTEM 1.0 (Grant et al., 2018). Using the cisTEM autorefine utility for 15 cycles and a reference obtained from the existing 5.5 Å map of *Lethocerus* thick filaments, EMD-3301, low pass filtered to 60 Å resolution, we obtained an initial ~8Å resolution map.

To further improve the map, we started from the beginning and extracted particles with a larger box size of 432 x 432 pixels, which is large enough to include ~ 5 crowns. Whole frame CTF correction was redone using GCTF (Zhang, 2016) with a particle count of 198,341. Later the whole frame CTF correction was replaced with a local CTF correction using GCTF. Using this enlarged data set we repeated 2D classification using *rome_map* and *rome_sml* and selected out 148,572 good segments.

We then returned to cisTEM to perform alignment without classification then repeated the 2D classification to remove more bad particles. Kept doing this cycle until there was little to no room for improvement and we got our latest map which is ~7Å. We used the earlier 8Å *Drosophila* reconstruction lowpass filtered to 50 Å for this reference. We found 3D classification not very helpful in the final reconstruction phase because we obtained little separation of segments into different classes and more than 87% percent of segments would go to one class. Finally, we rechecked the helical parameters using RELION 1.4 and we obtained 148.82 Å for the helical rise and +33.86° for the helical twist. Due to the magnification error, helical rise did not match the known axial rise of 145 Å which was resolved by rescaling the pixel size from 2.074 to 2.0207.

The major 3D reconstruction challenge was our inability to align segments without a starting reference. RELION 1.4 used initially was capable of enforcing helical symmetry and would have been the preferred method. However, with the myosin heads disordered we were unable to obtain an alignment to the 145 Å axial period. To help the alignment, we lowpass filtered the previously determined 3D structure of *Lethocerus* thick filaments to 65Å and used it as reference. At 65Å resolution, the non-myosin proteins are not individually resolved in the filament backbone. However, since they have a tendency to cluster between the levels of myosin heads, they may provide sufficient density variation along the filament backbone to align the 145 Å axial period. *Lethocerus* has the same axial rise as *Drosophila* according to X-ray fiber diffraction (Irving, 2006). The helical angle of *Drosophila* has not previously been reported. Switching to cisTEM for the reconstruction, the version of which was not capable of performing Iterative Helical Real Space Reconstruction (Egelman, 2000), meant sacrificing inclusion of the helical parameters during the reconstruction process. Nevertheless, we obtained the first 3D map using alignment to an initial reference while enforcing only C4 symmetry (Fig. 2C). Enforcing helical symmetry tended to smooth out the rod structure.

The rotational symmetry of *Drosophila* flight muscle thick filaments has not been reported so we reconstructed the two filament data sets using C2, C3 and C4 rotational symmetry. The C2 reconstruction appeared to have 4-fold symmetry, whereas the C3 reconstruction looked like a three bladed screw (Supplemental Fig. 3).

To determine the helical rise and twist in the cisTEM reconstruction, we used the *relion_helix_toolbox* in RELION and obtained a helical rise of 148.826 Å and helical twist 33.86°. The larger value compared to the axial repeat measured by X-ray fiber diffraction (Irving, 2006) is comparable to the difference observed for the *Lethocerus* thick filaments and attributable to magnification error. Then we moved to impose the helical symmetry on the map and compare it with the original 3D structure and other than it being smoother there seems to be no major difference between the two structures. For local resolution estimation we used Mono-res and last step was to sharpen the map using the local deblur embeded in Scipion.

The method used for the mutant data was very similar to what we have done before. Appion manual picker was used to pick filaments and 131,658 segments were extracted using Relion1.4 helical extraction. Using cisTEM 2D classification, we were left with 44,201 segments for 3D refinement which resulted in an ~8 Å resolution 3D structure. Helical twist for this structure is 33.92 and rise of 150.0 Å is reported using relion_helix_toolbox. By rescaling pixel size from 1.29 to 1.24 helical rise will be 145 Å which is the value we expect based on X-ray fiber diffraction.

### 6.5. Map Interpretation & Segmentation

An individual myosin molecule is 11 crowns in length or 1600 Å in length (11 x 145 Å. To obtain a density map of a single myosin rod structure as well as determine the rod arrangement within ribbons, required extending the reconstruction to 11-crowns in length, which was done utilizing the helical symmetry determined by RELION. Individual myosin rods and ribbons were segmented using UCSF Chimera. Because the myosin heads and the proximal S2 were disordered, we only observed the 10 crowns of rod tightly held within the thick filament backbone. After segmenting the rods, we were able to identify and isolate non-myosin proteins. Myosin heads were more challenging since they were barely visible at 8Å resolution. To see them we low pass filtered the map to 40 Å. As shown in (Fig. 3B,C), S2 is not resolved so the connection between the heads and the backbone is missing.

### 6.6. Quantification of Protein Stoichiometry: Liquid Chromatography Mass Spectrometry

Myofibril and thick filament samples were solubilized in 150 μl 0.1% RapiGest SF Surfactant (Waters Corporation) in 1.5 ml microcentrifuge tubes (50°C, 1 hour). Proteins were reduced by addition of 0.75 μl of 1M dithiothreitol to each tube and heating (100°C, 10 min). Cysteines were alkylated by addition of 22.5 μl of 100 mM iodoacetamide in 50 mM ammonium bicarbonate and incubation in the dark (22°C, 30 min). Proteins were digested to peptides by addition of 25 μl of 50 mM ammonium bicarbonate containing 5 μg of trypsin (Promega) and incubating (37°C, 18 hours). The samples were dried down in speed vacuum device. Trypsin was deactivated and RapiGest was cleaved by addition of 100 μl of 7% formic acid in 50 mM ammonium bicarbonate and heating (37°C, 1 hour). The resultant peptides were dried down. RapiGest was cleaved again by addition of 100 μl of 0.1% trifluoroacetic acid and heating (37°C, 1 hour). The resultant peptides were dried down and reconstituted a final time in 60 μl of 0.1% trifluoroacetic acid. The tubes were centrifuged at 18,800 RCF for 5 minutes (Thermo, Sorvall Legend Micro 21R) to pellet the surfactant. The top 57 μl of solution was transferred into a mass spectrometry (MS) analysis vial.

A 20 μl aliquot of each sample was injected onto an Acquity UPLC HSS T3 column (100 Å, 1.8 μm, 1 × 150 mm) (Waters Corporation) attached to a UltiMate 3000 ultra-high pressure liquid chromatography (UHPLC) system (Dionex). The peptides were separated and the UHPLC effluent was directly infused into a Q Exactive Hybrid Quadrupole-Orbitrap mass spectrometer (MS) through an electrospray ionization source (Thermo Fisher Scientific). Data were collected in data-dependent MS/MS mode with the top five most abundant ions being selected for fragmentation.

Peptides were identified from the resultant MS/MS spectra by searching against a drosophila proteome database downloaded from UniProt using SEQUEST in the Proteome Discoverer 2.2 (PD 2.2) software package (Thermo Fisher Scientific). The potential loss of methionine from the N-terminus of each protein (−131.20 Da), the loss of methionine with addition of acetylation (−89.16 Da), addition of carbamidomethyl (C; 57.02 Da), oxidation (M, P; 15.99 Da: M; 32.00 Da) and phosphorylation (S, T, Y; 79.98 Da) were accounted for by variable mass changes. The Minora Feature Detector (PD 2.2) was used to identify LC peaks with the exact mass, charge states, elution time, and isotope pattern as the SEQUEST derived peptide spectral matches (PSMs) across the samples in the entire study. Areas under each LC peak were calculated and reported in PD 2.2 as peptide abundances. Peptide abundances were exported to Excel (Microsoft). Relative protein abundances were estimated using a label-free approach which partially mitigates differences in each peptides ionization efficiency (O’Leary et al., 2019).

The average LC peak area of the top three peptides for myosin heavy chain, essential light chain and regulatory light chain and each thick filament associated protein being; flightin, myofilin, stretchin-MLCK, paramyosin mini-parmyosin, projectin, titin/kettin, and obscurin were determined in Excel. The relative abundance of myosin to each thick filament associated protein was determined from these average LC peak areas. Three separate relative abundances were determined for each myosin associated protein from the LC peaks of the myosin heavy chain and two myosin light chains and standard deviation were determined. Each myosin molecule is arranged as a dimer, consisting of two heavy chains and four light chains. With exception of paramyosin, which also exists as a dimer, each myosin associated protein exists as a monomer. To account for these structural arrangements, the relative abundances were divided by two, where appropriate and the number of myosin molecules per myosin associated protein ± standard deviation were reported.

## Supporting information

Daneshparvar et al.,Movie1

Daneshparvar et al.,Movie2

Daneshparvar et al.,Movie3

Daneshparvar et al.,Movie4

Daneshparvar et al.,Movie5

Daneshparvar et al.,Movie6

Supplemental Methods, Images and Movie Legends

## SUPPLEMENTAL INFORMATION

Supplemental information accompanies this paper …

## AUTHOR CONTRIBUTIONS

FA collected the image data, ND performed the reconstruction and wrote the paper; DWT prepared the specimen, HR contributed to the data processing, MJP and TO’L performed the mass spectroscopy analysis, KAT supervised the project and wrote the paper.

## ACKNOWLEDGEMENTS

This research was funded by NIH grant R01 GM30598 to KAT and R00 HL124041 to MJP. The Titan-Krios was partially funded by NIH grant S10 RR25081 to KAT. The DE-64 was purchased from funds provided by NIH grant U24 GM116788 to the SouthEast Consortium for Microscopy of MacroMolecular Machines. We thank Dr. Wu-Min Deng (FSU) and the members of his laboratory, particularly Yi-Chun (Jack) Huang for the gift of wild type, W1118, *Drosophila* flies and for maintaining our mutant fly stocks. We also thank Prof. James Vigoreaux (University of Vermont) for the gift of the Dmlc2^[Δ2-46; S66A,S67A]^ fly strain and Prof. Donald L. D. Caspar for his insightful comments throughout this process.

## DATA DEPOSITIONS

The reconstruction volume has been deposited in the EMDB under accession code EMD-0000. The raw frames and frame aligned images as well as metadata are deposited in EMPIAR under accession code 0000.

## References Cited

Ayer, G. & Vigoreaux, J. O. 2003. Flightin is a myosin rod binding protein. Cell Biochem Biophys, 38, 41–54.

Ayme-Southgate, A. & Southgate, R. 2006. Projectin, the elastic protein of the C-filaments. *In:* Vigoreaux, J. (ed.) Nature’s versatile engine insect flight muscle inside and out. Georgetown, TX: Landes Bioscience/Eurekah.com.

Barton, B. & Vigoreaux, J. O. 2006. Novel myosin associated proteins. *In:* Vigoreaux, J. O. (ed.) Nature’s versatile engine: insect flight muscle inside and out. Georgetown, TX: Landes Bioscience / Eurekah.com.

Becker, K. D., O’donnell, P. T., Heitz, J. M., Vito, M. & Bernstein, S. I. 1992. Analysis of Drosophila paramyosin: identification of a novel isoform which is restricted to a subset of adult muscles. J Cell Biol, 116, 669–81.

Beinbrech, G., Meller, U. & Sasse, W. 1985. Paramyosin content and thick filament structure in insect muscles. Cell Tissue Res, 241, 607–614.

Bullard, B., Burkart, C., Labeit, S. & Leonard, K. 2005. The function of elastic proteins in the oscillatory contraction of insect flight muscle. J Muscle Res Cell Motil, 26, 479–85.

Bullard, B., Leake, M. C. & Leonard, K. 2006. Some functions of proteins from the Drosophila sallimus (sls) gene. *In:* Vigoreaux, J. O. (ed.) Nature’s versatile engine: insect flight muscle inside and out. Georgetown, TX: Landes Bioscience / Eurekah.com.

Bullard, B., Luke, B. & Winkelman, L. 1973. The paramyosin of insect flight muscle. J Mol Biol, 75, 359–67.

Bullard, B. & Pastore, A. 2019. Through thick and thin: dual regulation of insect flight muscle and cardiac muscle compared. J Muscle Res Cell Motil.

Bullard, B., Sainsbury, G. & Miller, N. 1990. Digestion of proteins associated with the Z-disc by calpain. J Muscle Res Cell Motil, 11, 271–9.

Champagne, M. B., Edwards, K. A., Erickson, H. P. & Kiehart, D. P. 2000. Drosophila stretchin-MLCK is a novel member of the Titin/Myosin light chain kinase family. J Mol Biol, 300, 759–77.

Chew, M. W. & Squire, J. M. 1995. Packing of alpha-helical coiled-coil myosin rods in vertebrate muscle thick filaments. J Struct Biol, 115, 233–49.

Colegrave, M. & Peckham, M. 2014. Structural implications of beta-cardiac myosin heavy chain mutations in human disease. Anat Rec (Hoboken), 297, 1670–80.

Craig, R. 2017. Molecular structure of muscle filaments determined by electron microscopy. Appl Microsc, 47, 226–232.

Dickinson, M., Farman, G., Frye, M., Bekyarova, T., Gore, D., Maughan, D. & Irving, T. 2005. Molecular dynamics of cyclically contracting insect flight muscle in vivo. Nature, 433, 330–4.

Egelman, E. H. 2000. A robust algorithm for the reconstruction of helical filaments using single-particle methods. Ultramicroscopy, 85, 225–34.

Farman, G. P., Miller, M. S., Reedy, M. C., Soto-Adames, F. N., Vigoreaux, J. O., Maughan, D. W. & Irving, T. C. 2009. Phosphorylation and the N-terminal extension of the regulatory light chain help orient and align the myosin heads in Drosophila flight muscle. J Struct Biol, 168, 240–9.

Foth, B. J., Goedecke, M. C. & Soldati, D. 2006. New insights into myosin evolution and classification. Proc Natl Acad Sci U S A, 103, 3681–6.

Galkin, V. E., Vanloock, M. S., Orlova, A. & Egelman, E. H. 2002. A new internal mode in F-actin helps explain the remarkable evolutionary conservation of actin’s sequence and structure. Curr Biol, 12, 570–5.

Gasek, N. S., Nyland, L. R. & Vigoreaux, J. O. 2016. The Contributions of the Amino and Carboxy Terminal Domains of Flightin to the Biomechanical Properties of Drosophila Flight Muscle Thick Filaments. Biology (Basel), 5.

Grant, T., Rohou, A. & Grigorieff, N. 2018. cisTEM, user-friendly software for singleparticle image processing. Elife, 7.

He, S. & Scheres, S. H. W. 2017. Helical reconstruction in RELION. J Struct Biol, 198, 163–176.

Hu, D. H., Matsuno, A., Terakado, K., Matsuura, T., Kimura, S. & Maruyama, K. 1990. Projectin is an invertebrate connectin (titin): isolation from crayfish claw muscle and localization in crayfish claw muscle and insect flight muscle. J Muscle Res Cell Motil, 11, 497–511.

Hu, Z., Taylor, D. W., Reedy, M. K., Edwards, R. J. & Taylor, K. A. 2016. Structure of myosin filaments from relaxed Lethocerus flight muscle by cryo-EM at 6 Å resolution. Sci Adv, 2, e1600058.

Huxley, A. F. 1974. Muscular contraction. J Physiol, 243, 1–43.

Huxley, H. E. & Brown, W. 1967. The low-angle x-ray diagram of vertebrate striated muscle and its behaviour during contraction and rigor. J Mol Biol, 30, 383–434.

Irving, M. 2017. Regulation of Contraction by the Thick Filaments in Skeletal Muscle. Biophys J, 113, 2579–2594.

Irving, T. C. 2006. X-ray diffraction of indirect flight muscle from *Drosophila* in vivo. Nature’s Versatile Engine: Insect Flight Muscle Inside and Out. Springer.

Jung, H. S., Burgess, S. A., Billington, N., Colegrave, M., Patel, H., Chalovich, J. M., Chantler, P. D. & Knight, P. J. 2008. Conservation of the regulated structure of folded myosin 2 in species separated by at least 600 million years of independent evolution. Proc Natl Acad Sci U S A, 105, 6022–6.

Katzemich, A., Kreiskother, N., Alexandrovich, A., Elliott, C., Schock, F., Leonard, K., Sparrow, J. & Bullard, B. 2012. The function of the M-line protein obscurin in controlling the symmetry of the sarcomere in the flight muscle of Drosophila. J Cell Sci, 125, 3367–79.

Lakey, A., Ferguson, C., Labeit, S., Reedy, M., Larkins, A., Butcher, G., Leonard, K. & Bullard, B. 1990. Identification and localization of high molecular weight proteins in insect flight and leg muscle. EMBO J, 9, 3459–67.

Lakey, A., Labeit, S., Gautel, M., Ferguson, C., Barlow, D. P., Leonard, K. & Bullard, B. 1993. Kettin, a large modular protein in the Z-disc of insect muscles. EMBO J, 12, 2863–71.

Lee, K. H., Sulbaran, G., Yang, S., Mun, J. Y., Alamo, L., Pinto, A., Sato, O., Ikebe, M., Liu, X., Korn, E. D., Sarsoza, F., Bernstein, S. I., Padron, R. & Craig, R. 2018. Interacting-heads motif has been conserved as a mechanism of myosin II inhibition since before the origin of animals. Proc Natl Acad Sci U S A, 115, E1991–E2000.

Levine, R. J., Elfvin, M., Dewey, M. M. & Walcott, B. 1976. Paramyosin in invertebrate muscles. II. Content in relation to structure and function. J Cell Biol, 71, 273–279.

Levine, R. J., Kensler, R. W., Yang, Z., Stull, J. T. & Sweeney, H. L. 1996. Myosin light chain phosphorylation affects the structure of rabbit skeletal muscle thick filaments. Biophys J, 71, 898–907.

Levine, R. J., Kensler, R. W., Yang, Z. & Sweeney, H. L. 1995. Myosin regulatory light chain phosphorylation and the production of functionally significant changes in myosin head arrangement on striated muscle thick filaments. Biophys J, 68, 224S.

Linari, M., Brunello, E., Reconditi, M., Fusi, L., Caremani, M., Narayanan, T., Piazzesi, G., Lombardi, V. & Irving, M. 2015. Force generation by skeletal muscle is controlled by mechanosensing in myosin filaments. Nature, 528, 276–9.

Maroto, M., Arredondo, J., Goulding, D., Marco, R., Bullard, B. & Cervera, M. 1996. Drosophila paramyosin/miniparamyosin gene products show a large diversity in quantity, localization, and isoform pattern: a possible role in muscle maturation and function. J Cell Biol, 134, 81–92.

Mclachlan, A. D. & Karn, J. 1982. Periodic charge distributions in the myosin rod amino acid sequence match cross-bridge spacings in muscle. Nature, 299, 226–31.

Menetret, J. F., Schroder, R. R. & Hofmann, W. 1990. Cryo-electron microscopic studies of relaxed striated muscle thick filaments. J Muscle Res Cell Motil, 11, 1–11.

Misof, B., Liu, S., Meusemann, K., Peters, R. S., Donath, A., Mayer, C., Frandsen, P. B., Ware, J., Flouri, T., Beutel, R. G., Niehuis, O., Petersen, M., Izquierdo-Carrasco, F., Wappler, T., Rust, J., Aberer, A. J., Aspock, U., Aspock, H., Bartel, D., Blanke, A., Berger, S., Bohm, A., Buckley, T. R., Calcott, B., Chen, J., Friedrich, F., Fukui, M., Fujita, M., Greve, C., Grobe, P., Gu, S., Huang, Y., Jermiin, L. S., Kawahara, A. Y., Krogmann, L., Kubiak, M., Lanfear, R., Letsch, H., Li, Y., Li, Z., Li, J., Lu, H., Machida, R., Mashimo, Y., Kapli, P., Mckenna, D. D., Meng, G., Nakagaki, Y., Navarrete-Heredia, J. L., Ott, M., Ou, Y., Pass, G., Podsiadlowski, L., Pohl, H., Von Reumont, B. M., Schutte, K., Sekiya, K., Shimizu, S., Slipinski, A., Stamatakis, A., Song, W., Su, X., Szucsich, N. U., Tan, M., Tan, X., Tang, M., Tang, J., Timelthaler, G., Tomizuka, S., Trautwein, M., Tong, X., Uchifune, T., Walzl, M. G., Wiegmann, B. M., Wilbrandt, J., Wipfler, B., Wong, T. K., Wu, Q., Wu, G., Xie, Y., Yang, S., Yang, Q., Yeates, D. K., Yoshizawa, K., Zhang, Q., Zhang, R., Zhang, W., Zhang, Y., Zhao, J., Zhou, C., Zhou, L., Ziesmann, T., Zou, S., Li, Y., Xu, X., Zhang, Y., Yang, H., Wang, J., Wang, J., Kjer, K. M., et al. 2014. Phylogenomics resolves the timing and pattern of insect evolution. Science, 346, 763–7.

Morris, E. P., Squire, J. M. & Fuller, G. W. 1991. The 4-stranded helical arrangement of myosin heads on insect *(Lethocerus)* flight muscle thick filaments. J Struct Biol, 107, 237–249.

O’leary, T. S., Snyder, J., Sadayappan, S., Day, S. M. & Previs, M. J. 2019. MYBPC3 truncation mutations enhance actomyosin contractile mechanics in human hypertrophic cardiomyopathy. J Mol Cell Cardiol, 127, 165–173.

Patel, S. R. & Saide, J. D. 2005. Stretchin-klp, a novel Drosophila indirect flight muscle protein, has both myosin dependent and independent isoforms. J Muscle Res Cell Motil, 26, 213–24.

Pringle, J. W. S. 1978. The Croonian Lecture, 1977. Stretch activation of muscle: function and mechanism. Proc R Soc LondB Biol Sci, 201, 107–30.

Pringle, J. W. S. 1981. The Bidder Lecture, 1980 The Evolution of Fibrillar Muscle in Insects. J Exp Biol, 94, 1–14.

Qiu, F., Brendel, S., Cunha, P. M., Astola, N., Song, B., Furlong, E. E., Leonard, K. R. & Bullard, B. 2005. Myofilin, a protein in the thick filaments of insect muscle. J Cell Sci, 118, 1527–36.

Ramirez-Aportela, E., Vilas, J. L., Glukhova, A., Melero, R., Conesa, P., Martinez, M., Maluenda, D., Mota, J., Jimenez, A., Vargas, J., Marabini, R., Sexton, P. M., Carazo, J. M. & Sorzano, C. O. S. 2020. Automatic local resolution-based sharpening of cryo-EM maps. Bioinformatics, 36, 765–772.

Reedy, M. K. 1967. Cross-Bridges and Periods in Insect Flight Muscle. Am Zool, 7, 465–481.

Reedy, M. K., Leonard, K. R., Freeman, R. & Arad, T. 1981. Thick myofilament mass determination by electron scattering measurements with the scanning transmission electron microscope. J Muscle Res Cell Motil, 2, 45–64.

Sallan, L., Friedman, M., Sansom, R. S., Bird, C. M. & Sansom, I. J. 2018. The nearshore cradle of early vertebrate diversification. Science, 362, 460–464.

Scheres, S. H. 2012. A Bayesian view on cryo-EM structure determination. J Mol Biol, 415, 406–418.

Schmitz, H., Lucaveche, C., Reedy, M. K. & Taylor, K. A. 1994. Oblique section 3-D reconstruction of relaxed insect flight muscle reveals the cross-bridge lattice in helical registration. Biophys J, 67, 1620–33.

Squire, J. M. 1973. General model of myosin filament structure. 3. Molecular packing arrangements in myosin filaments. J Mol Biol, 77, 291–323.

Squire, J. M., Bekyarova, T., Farman, G., Gore, D., Rajkumar, G., Knupp, C., Lucaveche, C., Reedy, M. C., Reedy, M. K. & Irving, T. C. 2006. The myosin filament superlattice in the flight muscles of flies: A-band lattice optimisation for stretch-activation? J Mol Biol, 361, 823–38.

Swank, D. M., Vishnudas, V. K. & Maughan, D. W. 2006. An exceptionally fast actomyosin reaction powers insect flight muscle. Proc Natl Acad Sci US A, 103, 17543–7.

Taylor, K. A., Reedy, M. C., Cordova, L. & Reedy, M. K. 1984. Three-dimensional reconstruction of rigor insect flight muscle from tilted thin sections. Nature, 310, 285–91.

Taylor, K. C., Buvoli, M., Korkmaz, E. N., Buvoli, A., Zheng, Y., Heinze, N. T., Cui, Q., Leinwand, L. A. & Rayment, I. 2015. Skip residues modulate the structural properties of the myosin rod and guide thick filament assembly. Proc Natl Acad Sci USA, 112, E3806–15.

Toyoshima, C. 1989. On the use of holey grids in electron crystallography. Ultramicroscopy, 30, 439–444.

Vigoreaux, J. O., Saide, J. D., Valgeirsdottir, K. & Pardue, M. L. 1993. Flightin, a novel myofibrillar protein of Drosophila stretch-activated muscles. J Cell Biol, 121, 587–98.

Vinos, J., Domingo, A., Marco, R. & Cervera, M. 1991. Identification and characterization of Drosophila melanogaster paramyosin. J Mol Biol, 220, 687–700.

Wendt, T., Taylor, D., Trybus, K. M. & Taylor, K. 2001. Three-dimensional image reconstruction of dephosphorylated smooth muscle heavy meromyosin reveals asymmetry in the interaction between myosin heads and placement of subfragment 2. Proc Natl Acad Sci U S A, 98, 4361–6.

Wray, J. S. 1979. Structure of the backbone in myosin filaments of muscle. Nature, 277, 37–40.

Wu, J., Ma, Y. B., Congdon, C., Brett, B., Chen, S., Xu, Y., Ouyang, Q. & Mao, Y. 2017. Massively parallel unsupervised single-particle cryo-EM data clustering via statistical manifold learning. PLoS One, 12, e0182130.

Yang, S., Woodhead, J. L., Zhao, F. Q., Sulbaran, G. & Craig, R. 2016. An approach to improve the resolution of helical filaments with a large axial rise and flexible subunits. J Struct Biol, 193, 45–54.

Zhang, K. 2016. Gctf: Real-time CTF determination and correction. J Struct Biol, 193, 1–12.

