## Supplemental Methods, Images and Movie Legends for "CryoEM Structure of *Drosophila* Flight Muscle Thick Filaments at 7Å Resolution"

### Materials and Methods

#### Key Resources Table

| REAGENT or RESOURCE | SOURCE | IDENTIFIER |
|---|---|---|
| Chemicals, peptides, and recombinant proteins | | |
| Calcium insensitive gelsolin | M. Briggs, Duke University | |
| Talon Metal affinity resin | Clontech | #635503 |
| LB Media | Fluka | #L31562 |
| Carbenicillin | Sigma | #C1389 |
| Doxycycline | Sigma | #D9191 |
| IPTG | Sigma | #15502 |
| RapiGest SF Surfactant | Waters Corporation | |
| Porcine Erythrocyte Calpain | Athens Research & Technology, Athens, GA | # 16-05-030112-P |
| EXPERIMENTAL MODELS: ORGANISMS/STRAINS | | |
| *D. melanogaster* | Prof. Wu-Min Deng, Florida State Univ. | W1118 |
| *D. melanogaster* | Prof. J. Vigoreaux, Univ. Vermont | Dmlc2[Δ2-46; S66A,S67A] |
| *E.coli*: BL21(DE3) Competent cells | Novagen | |
| DEPOSITED DATA | | |
| *Lethocerus* thick filament | EMD-3301 | (Hu et al., 2016) |
| Titan Fibronectin3 repeat | PDB code 1TTG | (Main et al., 1992) |
| Titan Ig domain | PDB code 1TIT | (Improta et al., 1996) |
| Myosin Binding ProteinC | PDB code 2YXM | (Kishishita) |
| SOFTWARE AND ALGORITHMS | | |
| MotionCor2 | https://emcore.ucsf.edu/ucsf-motioncor2 | (Zheng et al., 2017) |
| CTFFIND4 | http://grigoriefflab.janelia.org/ctffind4 | (Rohou and Grigorieff, 2015) |
| GCTF | https://www.mrc-lmb.cam.ac.uk/kzhang/ | (Zhang, 2016) |
| RELION | https://www3.mrc-lmb.cam.ac.uk/relion//index.php/Main_Page | (He and Scheres, 2017) |
| cisTEM | https://cistem.org/ | (Grant et al., 2018) |
| EMAN2 | http://blake.bcm.tmc.edu/EMAN2/ | (Tang et al., 2007) |
| UCSF Chimera | https://www.cgl.ucsf.edu/chimera/ | (Pettersen et al., 2004) |

### Contact for Reagent and Resource Sharing

### ADDITIONAL METHOD DETAILS

#### Protein Expression and Purification

Calcium insensitive human plasma gelsolin (residues 25-406) was cloned and overexpressed in *Escherichia coli* BL21 (DE3) strain in Lysogeny Broth (LB) medium. The expression vector was obtained from Dr. Margaret Briggs (Duke University Medical Center). The protocol is as follows:

#### Buffers for Gelsolin preparation

1. LYSIS BUFFER (1X): 50 mM Tris, 0.1M (NaCl, 2mM EDTA with 0.1mM PMSF, 1mM DTT and 0.2mg/ ml lysozyme added, pH 7.9.
2. BINDING BUFFER (1X): 20 mM Tris, 50 mM NaCl, 5 Imidazole with 0.1 mM PMSF, and 6M Urea, pH 7.9.
3. WASH BUFFER (1X): 20 mM Tris, 50mM NaCl, 60 Imidazole + 6M Urea, pH 7.9
4. ELUTION BUFFER (1X): 20mM Tris, 50mM NaCl, 1M Imidazole + 6M Urea, pH 7.9
5. DIALYSIS BUFFER (1X): 10mM Mops, 20mM KCl, 5mM $MgCl_2$, 5mM EGTA, 0.5mM DTT, pH 6.8. Need 3 liters in total.

#### Gelsolin Preparation

Transformed cells were used to seed 80 ml of LB media + 0.1ml Carbinicillin and 50 µl Doxycycline. Culture was grown over night at 35° C and 242 rpm. The following day 30 ml of overnight culture is added to 500 ml LB + 0.5ml Carbinicillin and 1.0 ml Doxycycline and grown at 242 rpm and 35 °C for 3 hrs. The Optical Density (OD) is checked and gel samples made until an OD between 0.6 and 0.7 is reached at which point 400 µl of 1 M IPTG is added and gel samples at 1, 2, and 3 hours post IPTG.

At 3 hrs. the culture is centrifuged at 8K rpm, 4°C, for 15 min to pellet the cells. The supernatant is poured off and the pellet resuspended in PBS and transfered to a 50 ml centrifuge tube. After centrifuging at 7000 rpm, 4°C for 15 min, the pelleted cells are stored at -80°C.

For protein isolation, the pellet was thawed slowly on ice and suspended in 6 ml Lysis buffer +6 mg dry lysozyme. The thawed pellet was then shaken for 20 min at 250 rpm at 25°C.

Lysis buffer was then added to 25 ml and the suspension sonicated until large aggregates are no longer visible. The suspension, which consists primarily of gelsolin in inclusion bodies, was then transfer to 50 ml centrifuge tube and centrifuged at 11.5 rpm for 15 min. The pellet is resuspended in 20 ml Binding buffer plus 0.1 mM PMSF and 0.5% Triton and left for 20 min on ice. The suspension is pelleted by centrifugation at 20,300 x g for 15 min. Resuspended in binding buffer and repelleted. Finally, the pellet is resuspended in 5 ml Binding buffer + 6M Urea.

To purify the gelsolin, first charge a Talon column with about 70 ml 50 mM $CoCl_2$. Run 100 ml Binding buffer + 6M Urea.

Clarify the denatured gelsolin by centrifuging in a TLA100.3 rotor at 50,000 rpm for 30 min and load the supernatant on Talon column. Run 100 ml Binding buffer +10 mM Imidazole +6M Urea first to elute impurities followed by Binding buffer + 4 M imidazole + 6 M Urea. Take gel samples at regular intervals to check if gelsolin is not binding. After about 11 ml of 4M imidazole collect 2 ml fractions.

Read OD of fractions and save those fractions with the highest OD. Dialyze those in 1 liter Dialysis buffer with 3 changes. Centrifuge at 60,000x g, 30 min to clarify. Flash freeze small aliquots and store at -80°C.

**Buffers for Thick Filament Preparation**

1. RELAXING BUFFER (1X): 20 mM $Na_2HPO_4$, 80 mM KCl, 5 mM MgAcetate, 5 mM ATP, 5 mM EGTA, 1 mM DTT, pH 6.8.
2. CALPAIN BUFFER (1X): 10 mM Mops, 10 mM $Na_2HPO_4$, 80 mM KCl, 5 mM MgAcetate, 5 mM ATP, 5 mM EGTA, 3 mM DTT, 5.2 mM $CaCl_2$, pH 6.8.
3. SHEAR BUFFER (1X): 20 mM MOPS, 20 mM $Na_2HPO_4$, 100 mM NaCl, 5 mM MgAcetate, 5 mM ATP, 5 mM EGTA. 5 mM DTT, pH 6.8
4. STOP BUFFER (1X): 20 mM MOPS, 150 mM NaCl, 5 mM MgAcetate, 15 mM EGTA, pH 6.8.

Flight muscle from the thoraces of ~10 files is deposited in 0.3 ml of relaxing buffer +10 μl protease inhibitor (Sigma 2714). Muscle is then homogenized in a 1 ml ground glass homogenizer to a total volume of 1 ml (with rinses) and transfered to 1.5 ml centrifuge tube. Myofibrils are separated from solubilized proteins by centrifugation at 6,000x g with turns of the tube to promote pelleting. (3-4 turns, 3 min each). The pellet is resuspended in 0.3 ml of relaxing buffer +0.5% Triton and incubated on ice for 15-30 min. After incubation in relaxing buffer +Triton, the myofibril suspension is centrifuged at 6,000x g with tube turns (3-4 turns, 3 min each), resuspended in relaxing buffer and centrifuged again without the incubation time.

The pellet is then resuspended in 0.1 ml Calpain buffer +1 μl calpain (Athens Research). Digest at r.t. for 1 hour. Digest is stopped by adding 0.2 ml Stop buffer.

Digested myofibrils are then separated by centrifuging at 7,000x g, with turns (4 turns, 3 min each), the supernatant discarded, and 35-80 μl of Shear buffer depending on pellet size. Myofibrils are sheared 10X by pulling prep through a 1 ml syringe with 26G needle. Large solids are removed by centrifuging at 3,500x g with 2 turns (2 turns, 2 min each) to remove solids. The supernatant is collected and mixed with gelsolin. Typically 15 μl gelsolin (concentration 2 mg/ml) is used.

Thick filaments were checked quality and concentration by negative staining using 2% uranyl acetate. Large undigested material was removed by low speed centrifugation. Thick filaments are never subjected to high-speed centrifugation to separate them from intact or partially digested actin filament fragments or to concentrate them for EM grid preparation. If the thick filament suspension is too dilute, successive drops are deposited on the EM grid.

**Supplemental Figures**

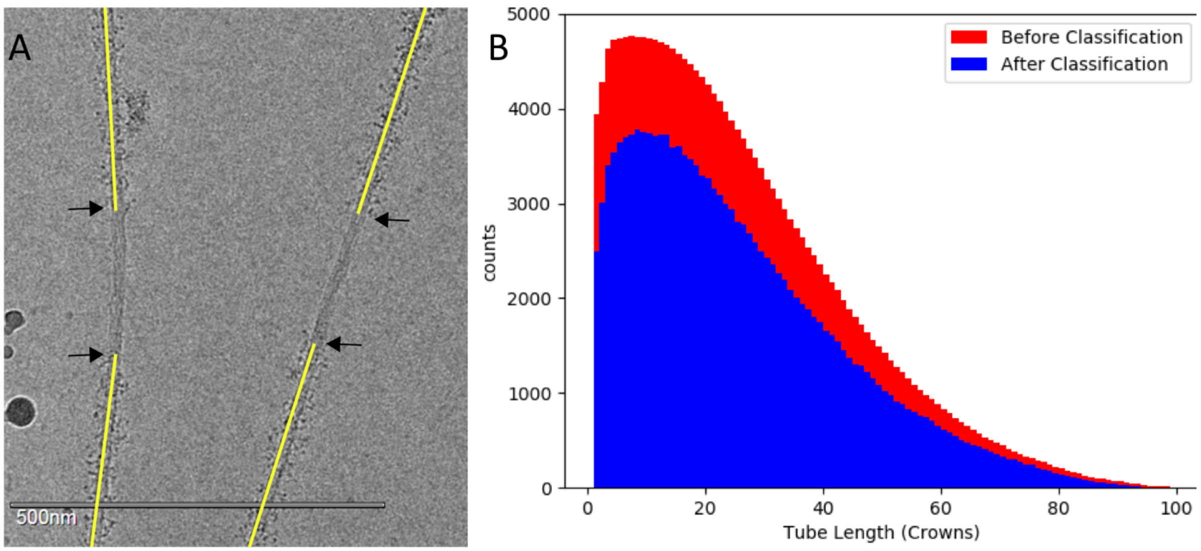

Figure S1. (A) Filaments are picked from the M-band (crown 0) but not all of them have the same length. Some of the filaments end outside of the image and others go over the carbon grid. Note that the disordered heads are clearly visible in the A-band region but the bare zone is very smooth. (B) Distribution of the tube length shows most segments used in the reconstruction come from an area close to the M-band. The distribution does not change after classification showing that distance from the M-band is not a major factor. *Drosophila* obscurin, the one protein whose visibility might be affected by a paucity of filament segments near the bare zone, is only long enough to reach the first crown of the A-band. The number of filament segments beyond 90 crowns is too small a fraction of the total for projectin to appear in the reconstruction.

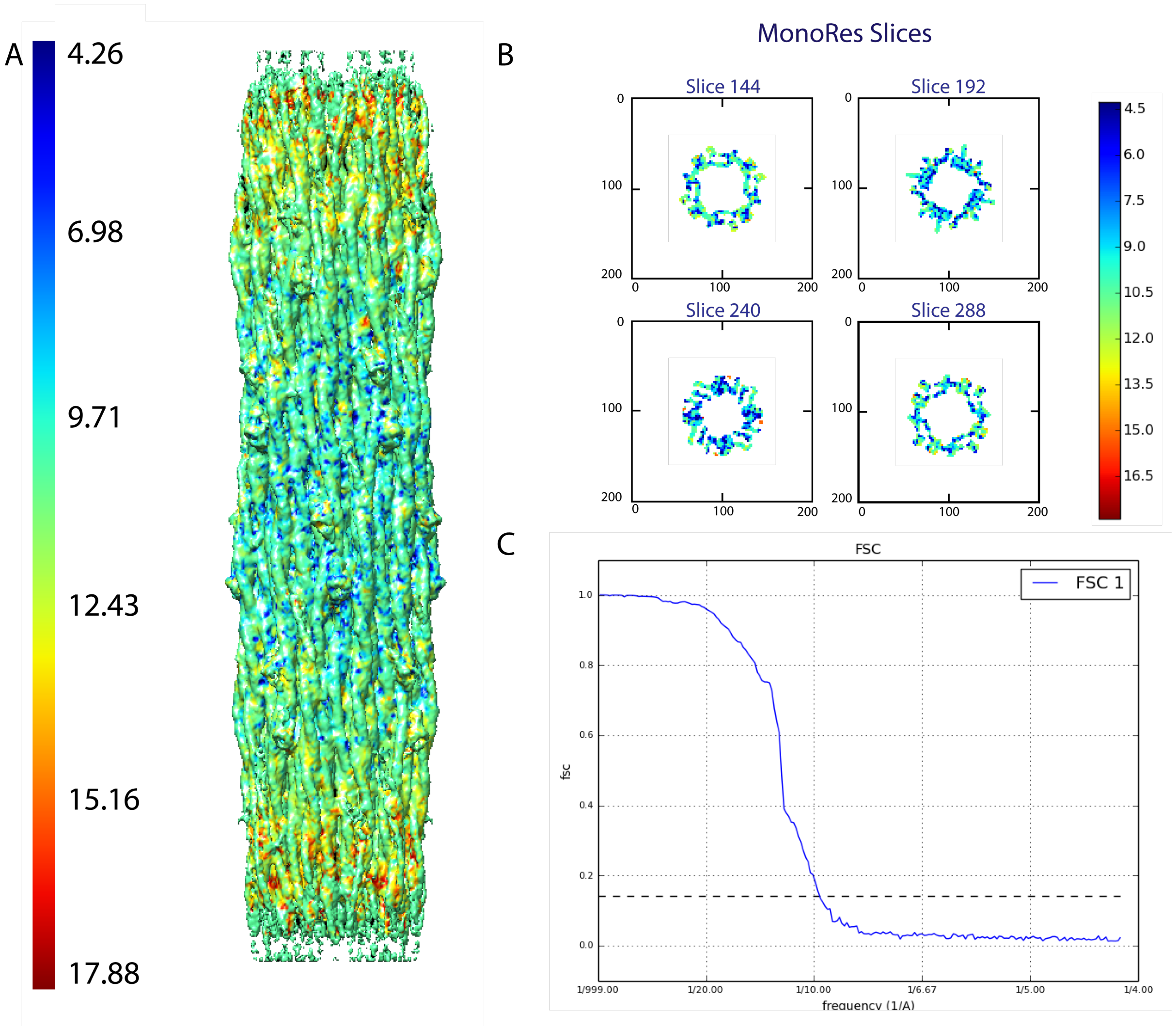

Figure S2. Resolution plots for the W1118 thick filament reconstruction computed by Monores embedded within Scipion (Gomez-Blanco et al., 2018). The color bars show the local resolution on each surface in Ångstroms. (A) Side view of local resolution map of the wild type Drosophila using. Map used was at 2.5 threshold which is the optimal threshold that includes most of the features with various local resolutions. Most of the map is at ~7.5Å resolution.. (B) Slices of the resolution map from in the Z direction. (C) FSC plot of the map generated from two half maps using Scipion.

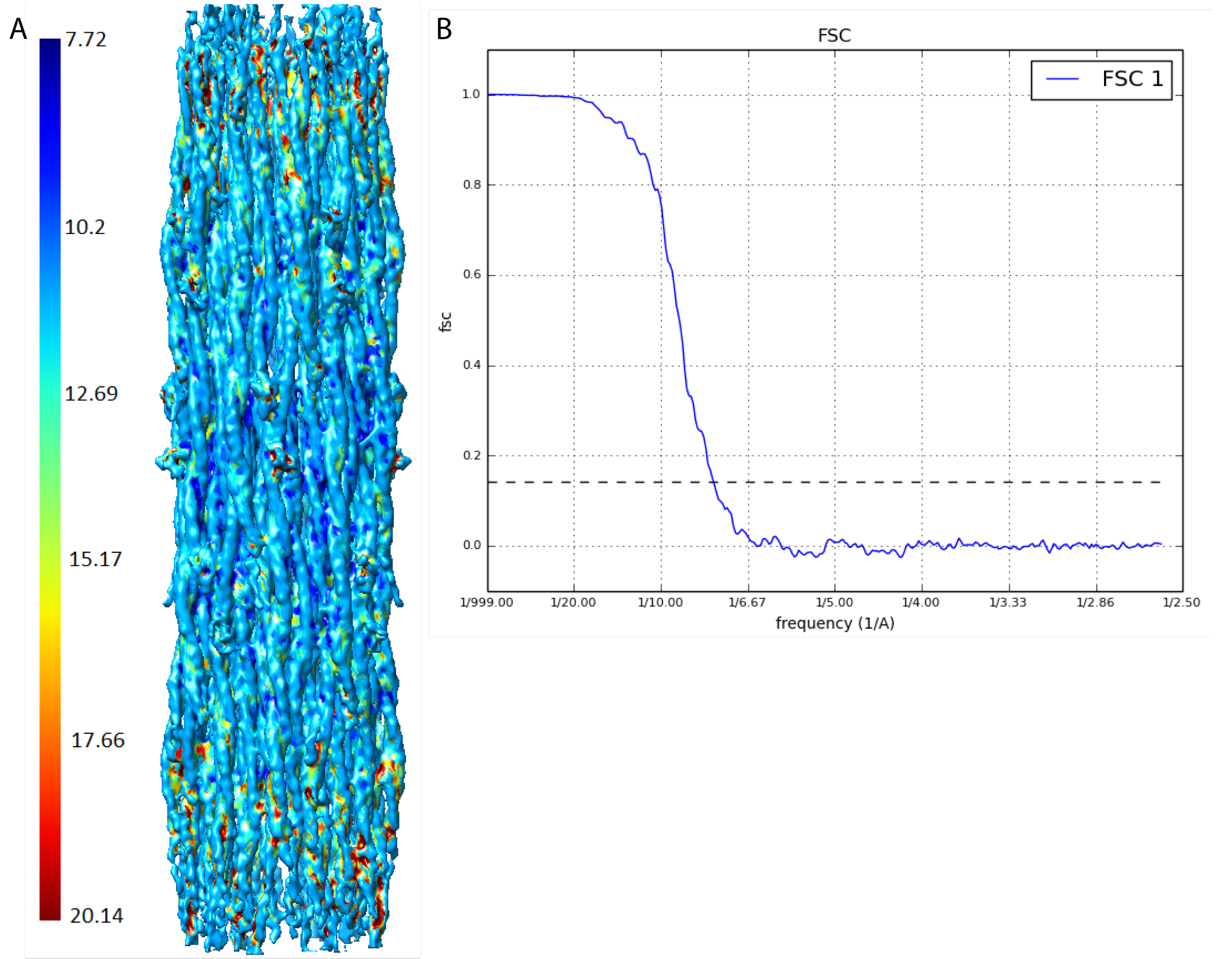

Figure S3. Resolution plots of the *Drosophila* Dmlc2[Δ2-46; S66A,S67A] mutant. The color bar shows the local resolution on the surface. (A) Side view. Map used was at 0.8 threshold which is the optimal threshold that includes most of the features with various local resolutions. Most of the map is at ~8 Å resolution. (B) FSC plot of the map generated from two half maps using Scipion.

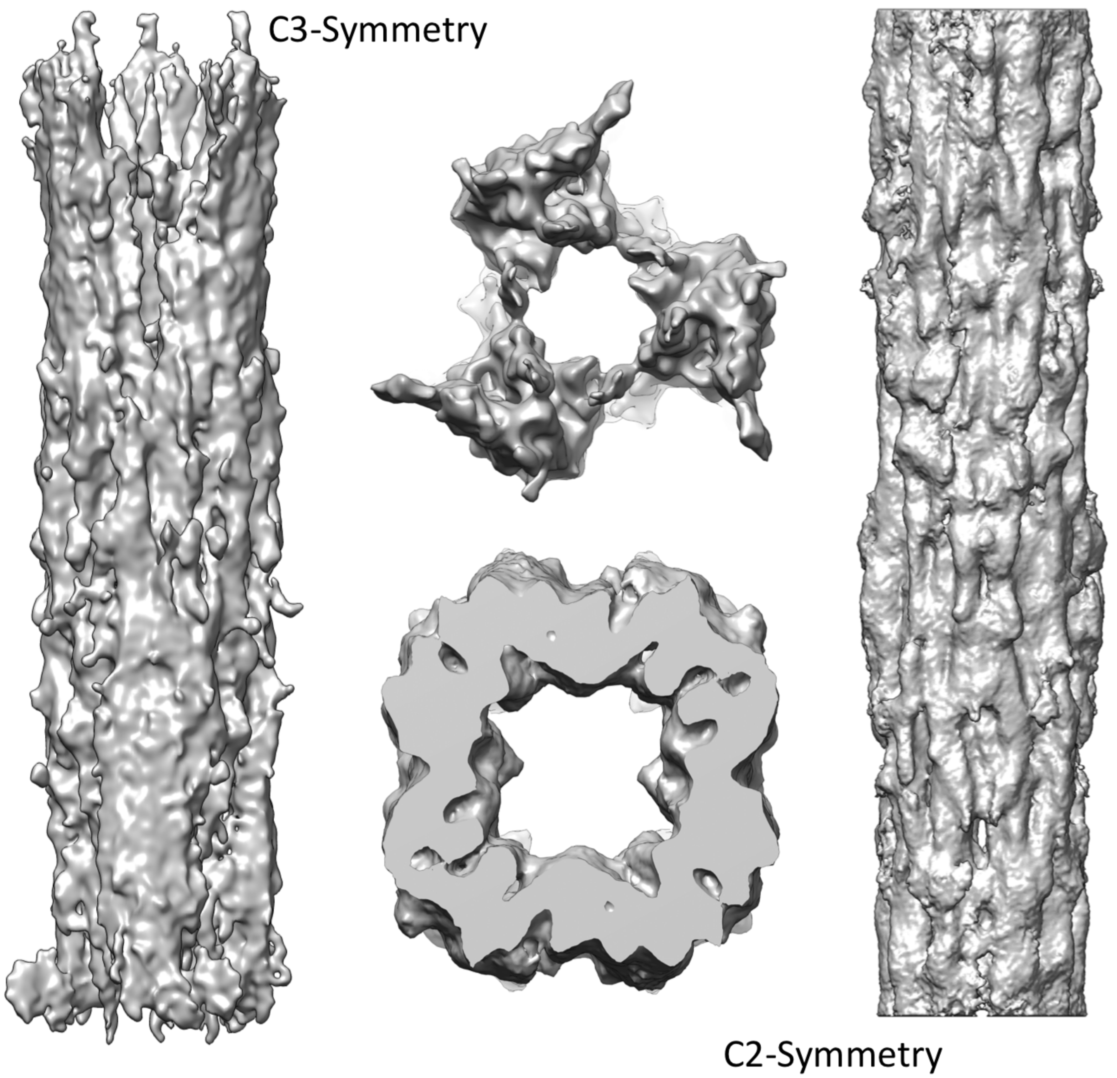

Figure S4. Reconstructions done using alternate symmetries. The reconstruction imposing C3 symmetry ( resulted in three rather massive densities with slim connections between them. The reconstruction done with C2 symmetry has more uniform density within which the myosin tails are arranged.

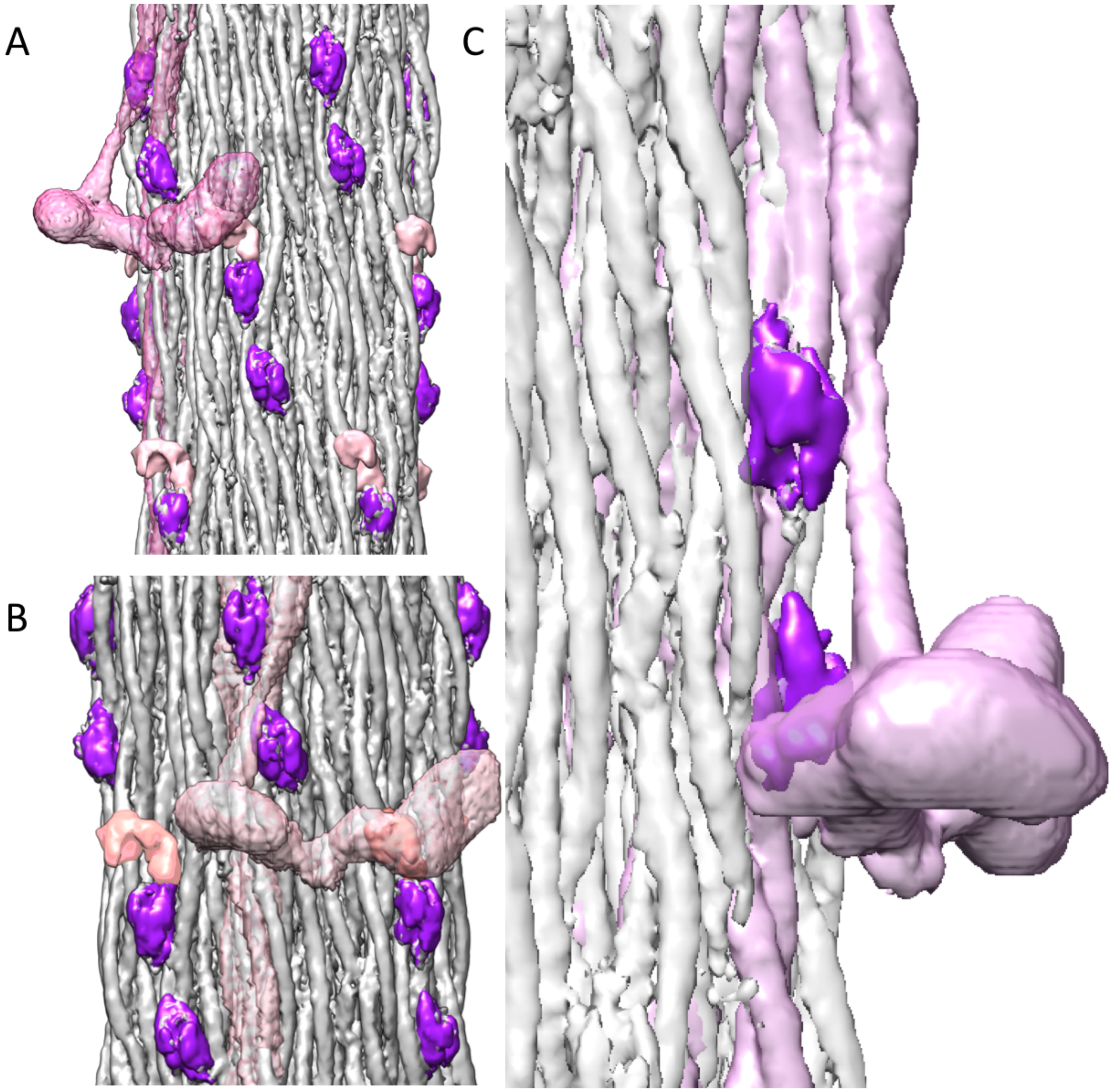

Figure S5. *Lethocerus* thick filament ribbon aligned to the Drosophila thick filament backbone with putative stretchin-klp Iset densities superimposed on a *Lethocerus* myosin ribbon with myosin heads. Iset domains are colored purple, *Lethocerus* myosin violet, *Drosophila* thick filament backbone is light grey and linker densities shown in pink. (A) Side view showing one Iset domain positioned near the proximal S2 of Lethocerus. (B) Front view showing one Iset domain and linker domain (pink) positioned near the free head RLC. (C) Side view from blocked head toward free head showing closeness of Iset domain to the proximal S2 and the myosin heads.

**Movie Legends**

Note that the orientation of all movies has the M-line at the top, Z-disk at the bottom.

**Movie 1**. Longitudinal slabs cutting through the segmented filament backbone. Myosin tails are colored in white, light grey and dark grey to differentiate the separate myosin ribbons. Flightin is colored red; myofilin, yellow; the purple and pink densities on the backbone surface are putative strechin-klp. The blue density is from an unknown non-myosin protein, possibly a domain of flightin or myofilin.

**Movie 2.** A 360° view of the filament backbone. Flightin is colored in red; purple and pink densities are putative strechin-klp. The proximal S2 connecting tails to myosin heads is not resolved due to the myosin heads being disordered. Orientation has the M-line at the top, Z-disk at the bottom.

**Movie 3**. The four main non-myosin densities observed in *Drosophila* flight muscle thick filaments. Flightin is red, myofilin yellow, putative stretchin-klp purple and pink, unknown blue. Three ribbons are colored drk gray, light gray and white. An I-set domain atomic structure (PDB 2YXM) displayed as a blue ribbon diagram has been fit into the putative stretchin-klp densities. The fit is good but not definitive at this resolution. Just before the movie starts, the myosin tail N-terminus, i.e. the beginning of the proximal S2 is visible in the upper right hand corner. Orientation has the M-line at the top, Z-disk at the bottom.

**Movie 4**. Putative strechin-klp Ig domains (purple) and long linker (pink) decorating the thick filament backbone. The pink colored feature we interpret as an average over several long linker structures. Consequently, its shape is possibly meaningless. The three densities appear to define a left-handed helical track, though the linkers between the features are not resolved. This is not a definitive assignment but seems more reasonable as the separation distance between the pink and purple densities is less with this interpretation. The "floating" densities are quite possibly the disordered myosin heads.

**Movie 5**. *Drosophila* ribbon (gray) superimposed and aligned with one from *Lethocerus*, shown in wire mesh. The movie shows the rather high similarity between the *Lethocerus* and *Drosophila* ribbons. The *Lethocerus* density shows the proximal S2 as well as a partial outline of the free head and the blocked head regulatory light chain.

**Movie 6.** Densities located on the thick filament backbone. At 50% transparency and aligned to the *Drosophila* backbone is a ribbon from the thick filaments from *Lethocerus indicus*. The contour cutoff for the *Lethocerus* ribbon is sufficient to show a partial outline of the free head and the blocked head regulatory light chain. Nevertheless, the *Lethocerus* myosin heads have enough volume to show close proximity to the stretchin-klp density (pink). One of the stretchin-klp densities also lies in close proximity to the *Lethocerus* proximal S2 where it leaves the close packing of the filament backbone.

**References Cited**

GOMEZ-BLANCO, J., DE LA ROSA-TREVIN, J. M., MARABINI, R., DEL CANO, L., JIMENEZ, A., MARTINEZ, M., MELERO, R., MAJTNER, T., MALUENDA, D., MOTA, J., RANCEL, Y., RAMIREZ-APORTELA, E., VILAS, J. L., CARRONI, M., FLEISCHMANN, S., LINDAHL, E., ASHTON, A. W., BASHAM, M., CLARE, D. K., SAVAGE, K., SIEBERT, C. A., SHAROV, G. G., SORZANO, C. O. S., CONESA, P. & CARAZO, J. M. 2018. Using Scipion for stream image processing at Cryo-EM facilities. *J Struct Biol,* 204**,** 457-463.

GRANT, T., ROHOU, A. & GRIGORIEFF, N. 2018. cisTEM, user-friendly software for single-particle image processing. *Elife,* 7.

HE, S. & SCHERES, S. H. W. 2017. Helical reconstruction in RELION. *J Struct Biol,* 198**,** 163-176.

HU, Z., TAYLOR, D. W., REEDY, M. K., EDWARDS, R. J. & TAYLOR, K. A. 2016. Structure of myosin filaments from relaxed Lethocerus flight muscle by cryo-EM at 6 Å resolution. *Sci Adv,* 2**,** e1600058.

IMPROTA, S., POLITOU, A. S. & PASTORE, A. 1996. Immunoglobulin-like modules from titin I-band: extensible components of muscle elasticity. *Structure,* 4**,** 323-37.

KISHISHITA, S., OHSAWA, N., MURAYAMA, K., TERADA, T., CHEN, L., LIU, Z., SHIROUZU, M., WANG, B., YOKOYAMA, S. Crystal structure of I-set domain of human Myosin Binding ProteinC. http://www.rcsb.org/structure/2YXM.

MAIN, A. L., HARVEY, T. S., BARON, M., BOYD, J. & CAMPBELL, I. D. 1992. The three-dimensional structure of the tenth type III module of fibronectin: an insight into RGD-mediated interactions. *Cell,* 71**,** 671-8.

PETTERSEN, E. F., GODDARD, T. D., HUANG, C. C., COUCH, G. S., GREENBLATT, D. M., MENG, E. C. & FERRIN, T. E. 2004. UCSF Chimera--a visualization system for exploratory research and analysis. *J Comput Chem,* 25**,** 1605-12.

ROHOU, A. & GRIGORIEFF, N. 2015. CTFFIND4: Fast and accurate defocus estimation from electron micrographs. *Journal of Structural Biology,* 192**,** 216-221.

TANG, G., PENG, L., BALDWIN, P. R., MANN, D. S., JIANG, W., REES, I. & LUDTKE, S. J. 2007. EMAN2: an extensible image processing suite for electron microscopy. *Journal of Structural Biology,* 157**,** 38-46.

ZHANG, K. 2016. Gctf: Real-time CTF determination and correction. *J Struct Biol,* 193**,** 1-12.

ZHENG, S. Q., PALOVCAK, E., ARMACHE, J. P., VERBA, K. A., CHENG, Y. & AGARD, D. A. 2017. MotionCor2: anisotropic correction of beam-induced motion for improved cryo-electron microscopy. *Nat Methods,* 14**,** 331-332.
